# Joint modeling of social genetic effects in mono- and pluri-specific groups: case study in intercrops

**DOI:** 10.64898/2026.03.27.714849

**Authors:** J. Salomon, J. Enjalbert, T. Flutre

**Affiliations:** Université Paris-Saclay, INRAE, CNRS, AgroParisTech, GQE, Le Moulon, Gif-sur-Yvette 91190, France

**Keywords:** plant-plant interactions, indirect genetic effects, mixing ability, phenotypic plasticity, incomplete design, agroecological mixtures, automatic differentiation on LMM

## Abstract

The genetics of interspecific groups remains largely unexplored, despite the central role of social (or indirect) genetic effects in shaping phenotypic expression within communities. Intercropping, i.e. the simultaneous cultivation of multiple crop species in the same field, offers a powerful model to harness these interspecific social effects. Such species mixtures provide well-documented agricultural benefits, yet few breeding frameworks have integrated the genetics of social interactions. Here, we address this gap by extending quantitative genetic theory to interspecific groups, with intercropping as a concrete and applied model case. We propose a quantitative genetic model that jointly analyzes intra and interspecific interactions within a unifying framework. Breeding values are decomposed into a direct component, shared in mono and mixed-crops, an interspecific social component corresponding to the effect of one species on another, and an intraspecific component that captures the social effects within a mono-genotypic stand of cloned plants. Statistically, this consists in simultaneously fitting several linear mixed models, one per stand type, all having direct breeding values in common. As no open-source software can fit such a complex mixed model, we provide such an implementation in C++ available in the R package plantmix. Simulations across various genetic (co)variance structures and sparse experimental designs showed accurate estimation of all genetic (co)variances and breeding values. With an incomplete, yet balanced design combining sole crops and intercrops, genetic gains in both systems were achievable simultaneously, enabling breeding strategies that progressively integrate intercropping into existing, sole-crop-only schemes. More broadly, this framework allows dissecting direct and social genetic effects when genotypes are observed in mono- and mixed-species situations, cultivated or not.

**Article summary:** The genetics of interspecific groups remains largely unexplored, despite the central role of social genetic effects in shaping phenotypic expression within communities. We address this gap by extending quantitative genetic theory to interspecific groups, with intercropping as an example. We propose a quantitative genetic model that jointly analyzes intra- and interspecific interactions within a unifying framework. Simulations showed accurate estimation of genetic (co)variances and breeding values. This framework supports breeding strategies where breeders want to gradually integrate pluri-specific trials in their mono-specific platforms, and allows dissecting direct and social genetic effects when genotypes are observed in mono- and mixed-species situations.

## Introduction

The phenotypic expression of complex traits is shaped not only by an organism’s genotype and its interactions with the physical environment but also by each and every biotic interaction experienced by the organism, with conspecifics or individuals from other species, be their pathogenic or symbiotic. The genetic determinants of these biotic interactions with the phenotypes of other individuals are called indirect genetic effects (Bijma, 2014; Griffing, 1967; Hamilton, 1964; Willham, 1963), also known as *social* genetic effects (Baud et al., 2017; Bijma et al., 2007; Frank, 2007). In groups and communities, the environment thus includes a social component shaped by the genotypes themselves — essentially, an environment made up of genes (Cheverud, 2003; Wolf, 2003). These social interactions impact the evolution and adaptation of populations and communities (Marjanovic et al., 2018; Moore et al., 1997; Santostefano et al., 2025; Wolf et al., 1998), with important consequences on the genetic response to selection when breeding animals (Bergsma et al., 2008; Craig & Muir, 1996; Muir, 2005; Sartori & Mantovani, 2013) or crops (Bančič et al., 2021; Griffing, 1976, 1977; Hamblin et al., 1976; Hill, 1990; Sampoux et al., 2020; Wright, 1985) on which we focus here. Indeed, crops present interesting specificities regarding social interactions, as modern breeding has been focussed on selecting best performing genotypes (varieties or cultivars) in monospecific and monogenotypic stands, a case rarely met in animals. Therefore social interactions of modern varieties have been shaped by this selective pressure for best performance at the level of the group composed of genetically identical individuals (Anten & Vermeulen, 2016; Bijma & Wade, 2008). This notably resulted in high-yielding genotypes in dense stands (Duvick, 2005; Perez et al., 2019; Reynolds et al., 1994), described as “commensal” genotypes (Donald, 1968).

But crops have also been traditionally cultivated in multispecific stands, such as the well known milpa (Vazeux-Blumental et al., 2024), with co-evolutionary consequences (Fréville et al., 2022). Such practices of intercropping, i.e growing two or more crop species simultaneously in the same field during a growing season (Ofori & Stern, 1987), are gaining interest, due to their ability at mitigating the risk of crop failure under adverse weather conditions (extreme events increasing with climate change) or in low-input cropping systems (X.-F. Li et al., 2021; Rao & Willey, 1980; Raseduzzaman & Jensen, 2017; Weih et al., 2021). Numerous studies have also demonstrated intercropping benefits in disease management (Boudreau, 2013; Zhu & Morel, 2019) and resource use efficiency, especially in cereal-legume mixtures which reduce dependence on nitrogen fertilizers (Bedoussac et al., 2014; Beillouin et al., 2021; Jensen et al., 2020; Li et al., 2023) . Despite these advantages, known for a (very) long time (Etheridge & Helm, 1924; Francis, 1978; Harwood, 2024; Ofori & Stern, 1987; Vandermeer, 1989), intercropping is still rarely practiced in high-GDP countries. One of the lock-in is the lack of varieties adapted to intercropping, as all breeding efforts focus on sole crops. Yet, intercropping in such countries increases in popularity, e.g., in organic agriculture (Yan et al., 2025). Therefore, reallocating part of the research investment to develop breeding strategies for intercropping (IC) alongside sole cropping (SC) is a worthy endeavour.

The necessity of a distinct breeding approach for intercropping arises from the potentially weak, or even lack of, correlation between varietal performances observed in sole cropping and intercropping systems, noted here r_G_ = cor(BV^SC^, BV^IC^), where ‘BV’ stands for breeding values (Annicchiarico et al., 2019; Haug et al., 2023). This resembles the classical framework of selecting genotypes across different environments, i.e. accounting for G×E interactions (Falconer, 1952), “interaction” meaning here “deviation from the additivity of the main G and E effects”. Here, it raises the question of indirectly selecting in sole cropping versus directly in intercropping, as quantified by the relative efficiency of indirect vs direct selection that depends on r_G_ but also on the square root of narrow-sense heritabilities in sole cropping and intercropping, noted h_SC_ and h_IC_ (Annicchiarico et al., 2019): RE := (h_SC_ / h_IC_) r_G_. A key distinction in crop mixtures compared to the classical case is that the environment includes a social component. A low correlation r_G_ is, at least partly, due to the plastic response of genotypes to the strong environmental shifts induced by intercropping, which introduces additional interactions with a second species, sometimes referred to as G×G interactions (Cheverud, 2003; Wolf, 2003), here again in the sense of statistical deviations.

To capture these complexities, quantitative genetics models the individual *i*’s phenotype in a binary mixture as the sum of its *direct* breeding value (DBV_i_, also called producer or direct genetic effect DGE), representing the additive effects of its genes on its own phenotype, and a *social* breeding value (SBV_j_, also called associate or indirect genetic effect IGE), representing the additive effects of the genes from its neighbor *j* on *i*’s phenotype (Griffing, 1967; Muir, 2005). As a result, the covariance between direct and social breeding values influences the direction of selection response. When this covariance is negative, as expected under competition for the same resources, selection based on DBV only can lead to negative genetic gain for the mixture as a whole, (Griffing, 1967; Muir, 2005). In this framework, the total breeding value of the focal individual depends both on its direct and social breeding values, which extends the classical breeding value concept (BV_i_^SC^) by incorporating heritable social effects (Bijma et al., 2007). The contribution of specific interactions between i and j, captured by the statistical deviations (DBV×SBV), are usually ignored, except in the studies of mixing ability. Indeed, the concept of mixing ability of genotypes *i* and *j* allows for evaluating their specific compatibility in mixtures, for mixtures, e.g., Wright 1985 for interspecific mixtures. In this case, where genotypes *i* and *j* belong to different species,the general mixing ability of the focal individual can be partitioned as GMA_i_^IC^ := DBV_i_ + SBV_i_^IC^, which refers to its average performance in mixture, and the specific mixing ability as SMA_ij_^IC^ := (DBV×SBV)_ij_^IC^ + (DBV×SBV)_ji_^IC^, which refers to the particular interactions of genotypes *i* and *j* mixed together. The general mixing ability, GMA_i_^IC^, will be noted BV_i_^IC^ from here on.

Several breeding models, both statistical and genetic, and strategies have been proposed to enhance intercropping breeding efficiency (Hamblin et al., 1976; Hill, 1990; Sampoux et al., 2020; Wright, 1985), usually with the goal of breeding both species for intercropping, but not always, e.g., with the usage of testers as in Moutier et al. (2022) and Dubey et al. (2024). Given the practical challenges of breeding for IC (e.g., combinatorial explosion), recent proposals include incomplete designs (Haug, 2021) and genomic prediction (Bančič et al., 2021). However these studies often focussed exclusively on IC without integrating SC varieties within the same program, hence considering independent breeding schemes, or a future where breeders have switched completely from SC to IC breeding. Such a complete split between breeding programs seems however unrealistic, in a market presently driven by sole cropping. Beyond such “either-or” scenarios, quantified by the relative efficiency above, we argue that a breeding strategy combining both SC and IC is needed. This calls for the development of genetic models jointly addressing intercropping and sole cropping data, in order to account for social interactions occurring in both cropping systems. Indeed, SC is not free of social interactions. This was notably the motivation for Forst et al. (2019) to integrate sole crop performance in the genetic analysis of cultivar mixtures. They introduced the specific mixing ability of the focal individual in sole cropping (SMA_ii_^SC^), which represents the intra-genotypic interactions in monovarietal stand, and hence contributes to its classical breeding value in SC so that BV_i_^SC^ = GMA_i_^SC^ + SMA_ii_^SC^. This approach hence enabled the joint analysis of sole-crop and cultivar mixtures, but it did not relate SMA_ii_^SC^ with DBV and SBV_i_^SC^, and it was restricted to varietal mixtures.

To address these issues, we hypothesized that i) jointly analyzing SC and IC data would improve the estimation of genetic (co)variances and values, and ii) incomplete, yet balanced designs with both cropping systems would provide accurate estimates at a reasonable experimental cost, eliminating the need for separate breeding programs and allowing efficient joint selection. Moreover, leveraging genomic relationships would further improve the accuracy of all estimates. To test these hypotheses, we aim to explore key questions: (i) which genetic model best supports joint SC and IC analysis, (ii) how combining SC and IC in experimental design influences parameter estimation accuracy, (iii) what are the accuracy gains when mobilizing genomic relationships, and (iv) what genetic gain can be expected for SC and IC under one generation of selection. Our approach involves conducting simulations based on parameter values derived from published field trials (Haug et al., 2023; Moutier et al., 2022) to evaluate the feasibility and effectiveness of joint breeding strategies.

## Materials and methods

Two species are considered here, a focal one (e.g., a cereal such as wheat, named hereafter Species 1), subject of the breeding effort, and a tester (e.g., a legume such as pea, Species 2). To investigate the potential of i) an incomplete design to estimate (co)variances and ii) a joint analysis of both sole crops (SC) and intercrops (IC), the materials and methods are subdivided into four sections: i) the genetic model used to decompose the genetic contributions to the phenotype (e.g., yield), ii) the statistical model and inference procedure, iii) the simulation scenarios, and iv) the selection procedure for both systems (SC and IC).

### Joint genetic modeling of yields in sole cropping and intercropping systems

We assumed a set of I genotypes (indexed by i) for Species 1 and a set of J genotypes (indexed by j) for Species 2. In each micro-plot, we also assumed that the trait of interest was phenotyped per species, e.g., grain yield, with grains sorted at harvest. All phenotypic values in intercropping were modeled using the following genetic equations:

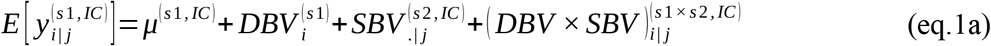

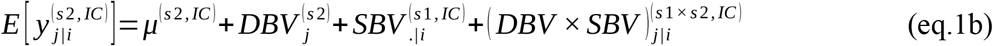

where 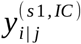 is the yield of the i-th genotype of species 1 mixed with the j-th genotype of species 2, indicated under the form “*i* | *j*” (read as “i given j”), *μ*^(*s*1, *IC* )^ is the mean yield of the species 1 in intercropping, 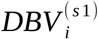 is the direct breeding value (DBV) of the i-th genotype of the species 1, 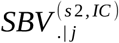 is the social breeding value (SBV) of the j-th genotype of species 2 on the yield of its partners from species 1, 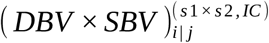 is the specific and orientated interaction where “*i* | *j*” indicates that it originates from genotype j and affects i-th phenotype; symmetrically for 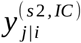.

In sole cropping, the phenotypic value can also be modeled with the direct breeding value but, in order to account for the genetic basis of phenotypic plasticity between sole crop and intercrop due to interactions between plants of the same genotype, we introduced a new breeding value, named the social intra-genotypic value (SIGV):

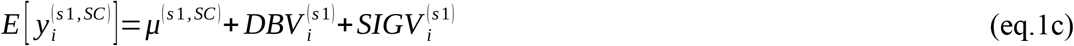

Similarly for the second species. The implication of such a genetic modeling is further detailed in the results section.

### Statistical modeling

We focused on trials in which the first species is the focal one whereas the second is made of testers, i.e., many more genotypes of the first species are observed compared to the second (I >> J), but the double factorial case is detailed in the Discussion. To estimate (co)variances and genetic values, three linear mixed models were defined based on the previous genetic model: i) equation 2 was used for the intercropping-only design, ii) equation 3 was applied for the sole crop-only design, and iii) both equations were jointly used for the designs with both sole crops and intercrops.

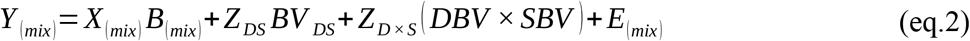

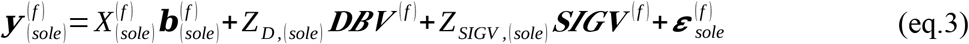

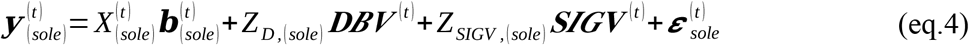

where *Y* _(*mix*)_ is matrix of yield observations in intercropping, with microplots in rows and one column per species, and **y** _(*sole*)_is the vector of yield observations from sole crops. *X*_(*mix*)_ and *X*_(*sole*)_ are the design matrices for the fixed effects in intercropping and sole cropping, respectively, such as species-specific intercepts and “block” effects as well as DBVs and SBVs from the tester species (which does not have enough levels to be modeled as random, but see the perspectives). *B*_(*mix*)_ is the matrix of fixed effects for intercropping, while ***b***_(*sole*)_ is the vector of fixed effects for sole crops. *Z*_*DS*_, *Z*_*D×S*_, *Z*_*D*_ and *Z*_*SIGV*_ are the design matrices for the random effects, i.e., BV_DS_ (the matrix with DBVs of the focal species in the first column and its SBVs in the second column), DBV×SBV (the matrix with (DBV×SBV)_i|j_ in the first column and (DBV×SBV)_j|i_ in the second column), DBVs only from the focal species and its SIGVs, respectively. *E*_(*mix*)_ and ***ε*** _(*sole*)_ denote the matrix and vectors of residual errors for IC and SC, respectively.

The DBVs and SBVs from the focal species were assumed to follow a matrix-variate Normal distribution (Furlotte & Eskin, 2015):

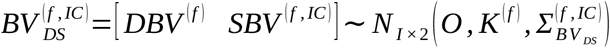

in which 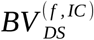 is a I×2 matrix of DBVs and SBVs, with I the number of focal genotypes, *O* is the mean matrix of zeros, *K*^(*f*)^ is the covariance matrix between the rows (kinship), corresponding to the additive genetic relationships between focal genotypes, and 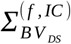 is the 2×2 covariance matrix between DBVs and SBVs, such that:

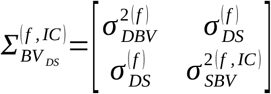

where 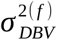 is the direct additive genetic variance, 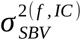 is the social additive genetic variance, and 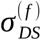 is the direct-social additive genetic covariance (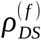 denoting the correlation).

The (DBV×SBV)^IC^ were assumed to follow a matrix-variate Normal distribution:

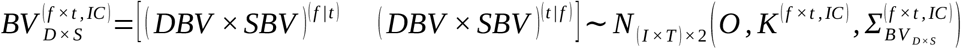

in which 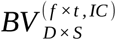 is a matrix of I×T rows and 2 columns, with I the number of focal genotypes and T the number of tester genotypes, *O* is the mean matrix of zeros, *K*^(*f ×t, IC*)^ is the covariance matrix between the rows, corresponding to the genetic similarity between mixtures (see the Discussion), and 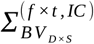 is the 2×2 covariance matrix between columns, such that:

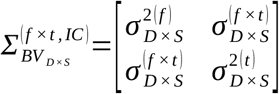

Moreover, for the focal species, 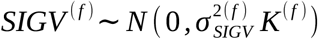, with 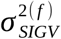 the social intragenotypic additive genetic variance whereas *SIGV* ^(*t*)^ is the vector of social intra-genotypic values of the tester species, modeled as fixed.

The residual errors in IC were assumed to follow a matrix-variate Normal distribution:

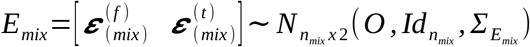

in which *E*_*mix*_ is a *n*_*mix*_× 2 matrix of errors within the same plots in intercropping, *n*_*mix*_ is the number of unique mixed plots, *O* is the mean matrix of zeros, 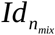 is the covariance between the rows (as an identity matrix), and 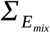 is the 2×2 covariance matrix between the columns, such that:

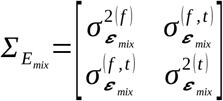

where 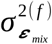 (respectively, 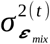) is the error variance for wheat (resp. pea).

In SC, the error term for focal and tester species was assumed to be distributed as 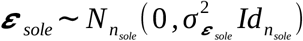 with *n*_*sole*_ the number of sole plots, and 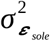 the error variance for sole plots and 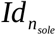 the identity matrix, respectively.

### Statistical inference and software implementation

Existing statistical tools in R, such as lme4, MM4LMM, ASReml-R or sommer, do not have the flexibility to fit correlated random effects (DBVs and SBVs) while also allowing the user to provide both a matrix for IC phenotypes and a vector for SC phenotypes as response variables. Custom code was hence developed to fit the new model in a computationally efficient way. The inference was conducted using Restricted Maximum Likelihood (REML) with Automatic Differentiation (AD) and Laplace Approximation (LA) via the Template Model Builder (TMB) R package (Kristensen et al., 2016; R Core Team, 2021). TMB was chosen for its open-source nature, promoting reproducibility, its flexibility in handling complex random-effects models, and its ability to produce accurate estimates efficiently compared to standard software used in the research community. By leveraging AD, TMB efficiently computes notably the first-order derivative of any objective function to optimize, here the negative log-likelihood, minimizing numerical instability, and reducing the computational burden by leveraging data structures tailored for sparse matrices. The Laplace Approximation in TMB, when integrating the random effects (and the fixed effects in REML), further enhances computational efficiency, and enables robust and scalable inference for high-dimensional mixed-effects models.

Our implementation was benchmarked to the ASReml-R reference software (The VSNi Team, 2023), using a single simulated dataset with the sole_only and inter_only designs (Table 1 and Figure 4 in File S2) to assess accuracy and computational efficiency. A laptop with 16 GB of RAM, CPU max MHz of 3700, running on the Debian GNU/Linux 13.2 operating system, was used for all analyzes.

### Simulation of a genomic relationship matrix

A panel was simulated, consisting of I=200 genotypes for the focal species (s1) and J=2 genotypes for the tester species (s2). SNP genotypes were simulated for the focal species using the sequential coalescent with recombination (Staab et al., 2015). Ten distinct chromosome pairs were considered, each 10^5^ bp in length. The mutation rate and recombination rate were both set to 10^−8^, the effective population size (Ne) to 10^4^ and migration rates between sub-populations to 10, so that the genetic structure is low. The simulation was implemented using the scrm R package, wrapped within the rutilstimflutre R package (Flutre, 2019). The resulting patterns of linkage disequilibrium (LD) and genetic structure are presented in (Figure 1 and Figure 3 in File S2). The additive genomic relationship matrix (GRM) was computed using the NOIA method (Vitezica et al., 2017) as implemented in the estimGenRel function of rutilstimflutre (R package).

**Figure 1 caption:**
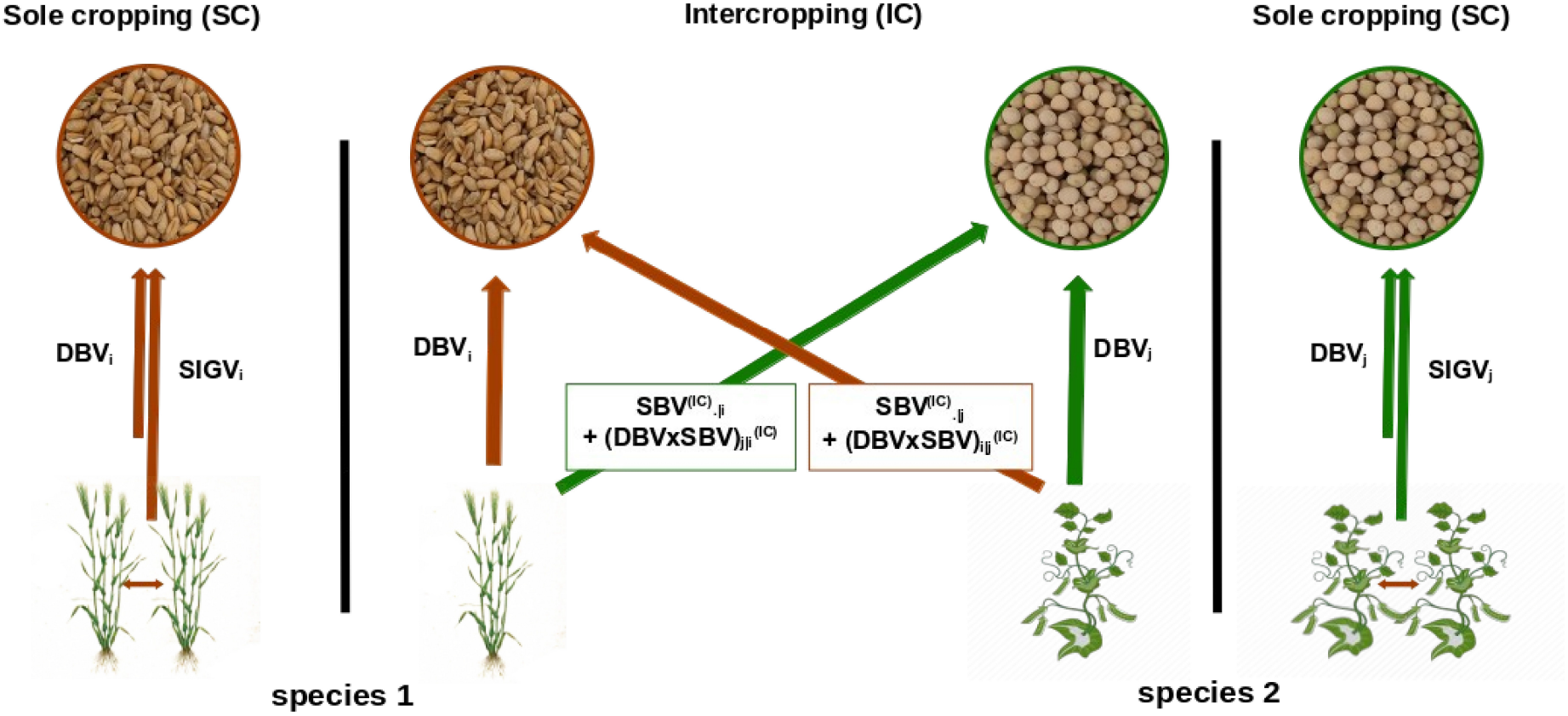
The proposed modeling framework of DBVs (direct breeding values), SBVs (social breeding values), DBV×SBVs (interactions between DBVs and SBVs) and SIGVs (social intragenotypic values) in the case of an interspecific binary mixture and both corresponding sole crops. The colors of the arrows and boxes correspond to the species trait affected by the given breeding values (e.g., the sum SBV _.|i_ + (DBVxSBV)_j|i_ originates from species 1 and affects species 2, it hence is colored in green).

### Simulations of field trials

#### Experimental designs

Three experimental designs were evaluated, each with a total of 400 micro-plots. In sole_only, the 200 focal genotypes are observed in sole cropping using a randomized complete block design with two blocks. In inter_only, all plots are observed in intercropping, according to a sparse design in which half focal genotypes were combined with the first tester and the remaining half with the second tester, both groups being evenly randomized across both blocks. In sole_inter*_*50, all focal genotypes are evaluated with no replication in sole cropping while half focal genotypes are tested with the first tester and the other half focal genotypes with the second tester, all plot types being evenly randomized across both blocks.

#### Parameters

The mean yield of each species in sole crop and intercrop was based on experimental data of wheat–pea field trials (Moutier et al., 2022): X (respectively, Y) for the focal species in solecrop (resp., intercrop), and X (resp., Y) for the tester species in sole crop (resp., intercrop). The genetic coefficient of variation for the DBV of the focal species was set to 8%, leading to a direct genetic variance 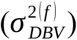 of 27.04. Five levels were considered for both the social genetic variance 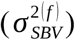 and the social intragenotypic variance 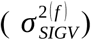 of the focal species: 10%, 20%, 30%, 40%, and 50% of the direct genetic variance (combinations of parameter values are hereafter called “scenarios”). The DBV–SBV correlation for the focal species was obtained based on experimental data of barley-pea field trials (Haug et al., 2023), i.e., set to -0.8 (see **File S1**). For each species s, the narrow-sense heritability in sole cropping (h^2^_SC,s_) and intercropping (h^2^_IC,s_) were both set to 0.7. The corresponding error variances were approximated using the following formulas: 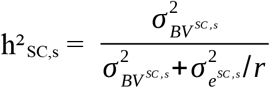 and 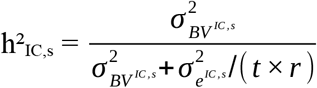, with *s* the species (focal or tester), *r* the number of blocks (2 here), and *t* the number of testers with which each genotype is observed (1 for inter_only). For the sole_inter_50 design, the same error variances were used (but see the Discussion). Overall simulation scenarios, the correlations between BV^SC^ and BV^IC^ ranged from 0.92 to 0.58.

Interspecific SMAs could also occur in intercropping, but were ignored in the simulations shown here, due to evidence for them often being small or absent (e.g., Haug et al., 2023, Salomon et al, in prep). Due to the low number of simulated testers (two), their effects were statistically modeled as fixed.

### Evaluation of estimation accuracy

For each scenario, one hundred datasets (“replicates”) were simulated. The statistical model above was then fitted to each of them. Estimation accuracy was assessed both at the level of the (co)variance parameters as well as the genetic values. For each genetic variance 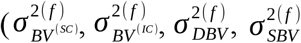 and 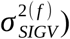 and correlation 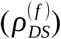, the bias error between the estimated and true value was computed per replicate i (BE_i_ := estim_i_ - true_i_), then normalized using the true value (NBE_i_ := BE_i_ / true_i_). To test that the mean NBE was higher than 0.15 in absolute value, two unilateral t tests were made. The null hypothesis was rejected when at least one p value was below 0.025. For each kind of genetic values (BV^SC^, BV^IC^, DBV, SBV and SIGV), the Pearson correlation per replicate i was computed between the empirical BLUPs and the true values. For the sole_only design, the breeding values in intercropping (BV^IC^) and their variance were approximated by the classical breeding values (BV^SC^) and their variance: 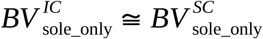 . Conversely, for the inter_only design, the breeding values in sole cropping (BV^SC^) and their variance were approximated by the direct breeding value (DBV) and their variance: 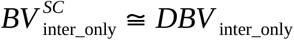.

### Evaluation of selection accuracy

All experimental designs were evaluated based on their ability to select the best genotypes of the focal species for sole cropping (highest BV^SC^) and for intercropping (highest BV^IC^) using the known simulated values. For the sole cropping target, genotypes were selected based on their estimated BV^SC^ in the sole_only design, on their estimated DBV in the inter_only design, and on the sum of their estimated DBV and SIGV in the sole_inter_50 design. For the intercropping target, genotypes were selected based on their estimated BV^SC^ in the sole_only design, and on the sum of their estimated DBV and SBV in the inter_only and sole_inter_50 designs. The true positive rate (TPR) was computed as the proportion of selected genotypes based on estimated breeding values that were matching the truly best genotypes based on their true breeding values (known here because they were simulated). A selection intensity of 10% was applied to select the best genotypes.

Specifically for the sole_inter_50 design, we introduced a new selection index:

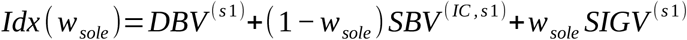

with *w*_*sole*_ weighing how much of the selection should be targeted towards sole cropping rather than intercropping. Its extreme cases correspond to the selection index for sole cropping only (*Idx* (1)=*Idx* _*SC*_ =*DBV* ^(*s*1)^+ *SIGV* ^(*s*1)^) and for intercropping only (*Idx* (0)=*Idx*_*IC*_ =*DBV* ^(*s*1)^+ *SBV* ^(*IC, s*1)^). Here the selection index was computed for each genotype with *w*_*sole*_ in {0.25, 0.5, 0.75}.

## Results

### Joint genetic modeling of sole and intercrops

A genotype i of species s_1_ intercropped with a genotype j of species s_2_ has both a direct and a social breeding value, noted DBV_i_ and SBV_i_^IC^, in lieu of a single breeding value in sole crop, BV_i_^SC^. To benefit from the joint analysis of both sole and intercrop data, it is necessary however to have one breeding value explaining part of the phenotypic variance in both crop types, so that both data sources can contribute to the estimation of the breeding value in question. We propose here to achieve this by expressing BV_i_^SC^ as the sum of DBV_i_ and of another term named SIGV_i_ for social intra-genotypic value. This was inspired by earlier work on varietal mixtures, and SIGV_i_ can itself be decomposed into more fundamental breeding values operating in “monovarietal” stands (shorten in “monovarietals”, also called “pure stands”), both of which justifies a detour.

In the context of monovarietals and varietal mixtures for which only the total yield is available, Forst et al.(2019) mobilized the general and specific mixing abilities, noted GMA_i_^SC^ and SMA_ij_^SC^ when the i-th genotype is mixed with the j-th genotype. The “SC” exponent indicates that both genotypes belong to the same species. Assuming equal plant densities, the total yield of such a binary mixture can hence be decomposed into ½GMA_i_^SC^ + ½ GMA_j_^SC^ + SMA_ij_^SC^. Most importantly, Forst et al. (2019) introduced the notion of “intra-genotypic interaction” (SMA_ii_^SC^) to capture how the i-th genotype performs in monovarietal compared to the mean of varietal mixtures comprising it. In this model, the classical breeding value in sole crop is defined as: BV_i_^SC^ := GMA_i_^SC^ + SMA_ii_^SC^.

In the intercropping case that is the focus here, where the i and j genotypes belong to two different species, the general and specific mixing abilities can be noted as GMA_i_^IC^, GMA_j_^IC^ and SMA_ij_^IC^. Moreover, it is more common in an intercrop to have access to phenotypic data *per species component*, allowing to define these mixing abilities directly in terms of direct and social breeding values (e.g., Sampoux et al., 2020; Wright, 1985): GMA_i_^IC^ := DBV_i_ + SBV_i_^IC^, and SMA_ij_^IC^ := (DBV×SBV)_ij_^IC^ + (DBV×SBV)_ji_^IC^. The notion of “breeding value in intercropping” can be defined as BV_i_^IC^ := GMA_i_^IC^ (e.g., as used in Bančič et al., 2021). Assuming the availability of phenotypic data at the scale of individual genotype components in a varietal mixture hence yields an equivalent expression, e.g., for the phenotype of genotype i from species s_1_ when mixed with genotype j from the same species: ½[DBV_i_+SBV_i_^SC^+(DBV×SBV)_ij_^SC^]. Note that SBV_i_^SC^ ≠ SBV_i_^IC^. Applying it to the classical breeding value in sole crop gives: BV_i_^SC^ := DBV_i_ + SBV_i_^SC^ + (DBV×SBV)_ii_^SC^.

When, as in our case here, only sole crop (i.e., monovarietal) and intercrop data are available, i.e., in the absence of additional data from varietal mixtures, the terms SBV_i_^SC^ and (DBV×SBV) _ii_^SC^ are not separately identifiable, but their sum is, which we named “social intra-genotypic value”: SIGV_i_ := SBV_i_^SC^ + (DBV×SBV)_ii_^SC^ = SBV_i_^SC^ + SMA_ii_, so that BV_i_^SC^ := DBV_i_ + SIGV _i_.

In brief, we introduced here a general framework decomposing performances in sole crop and intercrops in terms of all these breeding values. We also propose to clarify the orientation of the social genetic effects by introducing a new notation “a|b”, where “a” indicates the genotype(s) receiving the effect and “b” corresponds to the genotype at the origin of the effect. As a result, SBV _i_ is replaced by SBV_.|i_ corresponding to the social breeding value of genotype i on any other genotype it is cultivated with, and (DBV×SBV)_ij_ is replaced by (DBV×SBV)_j|i_ corresponding to the specific and orientated interaction in the i-j mixture, originating from the genotype i and having an effect on genotype j. Figure 1 illustrates this modeling framework as applied to cereal (species 1)–legume (species 2) intercrops and the respective sole crops.

### Accuracy of (co)variance estimates

To explore this new genetic model, three experimental designs were compared, all comprising the same total number of microplots. The sole_only design only involved sole crops and the inter_only design only intercrops, while the sole_inter_50 involved both sole and intercrops (Figure 2).

**Figure 2 caption:**
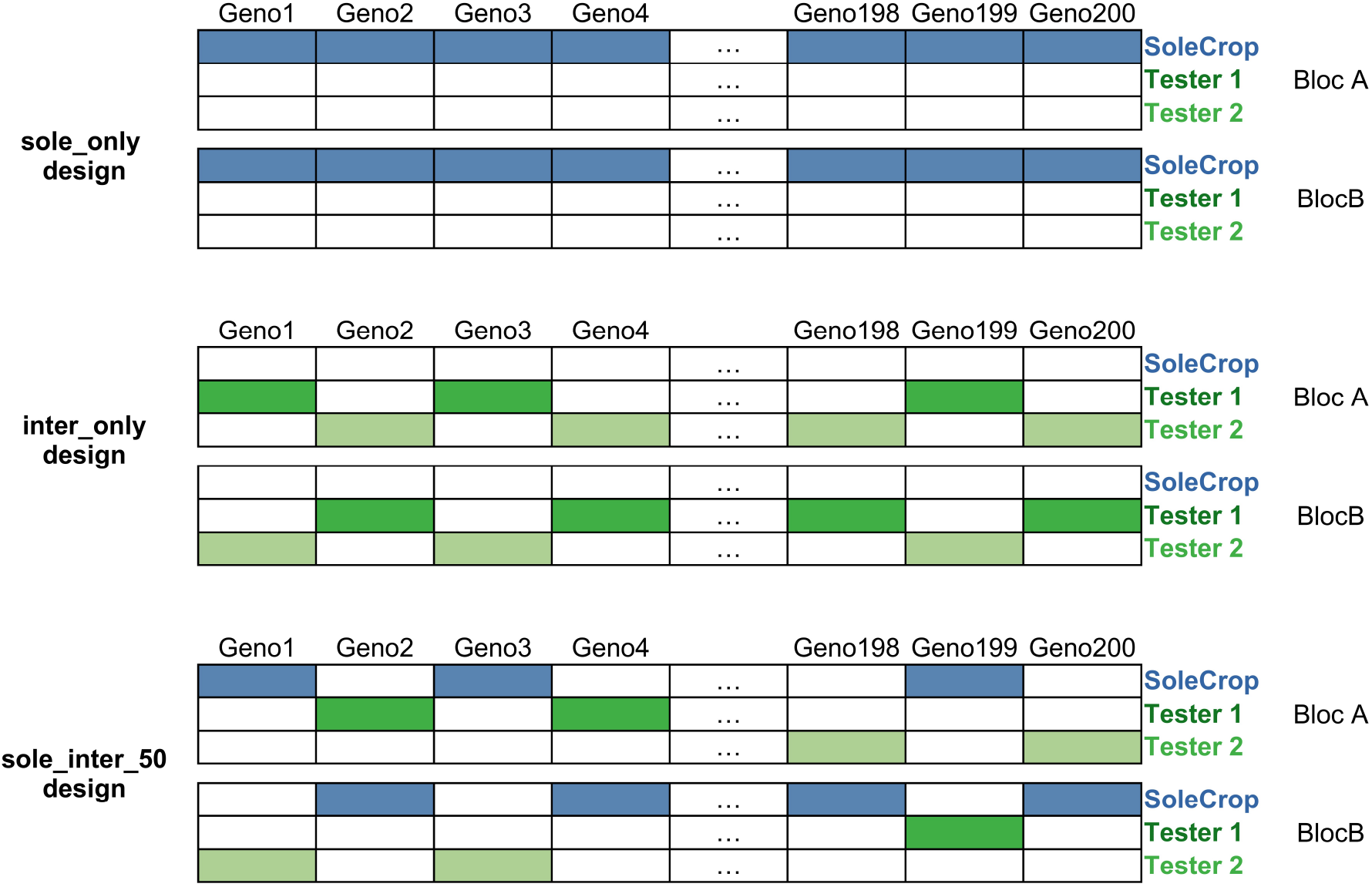
The three experimental designs compared in this study, each having the same number of plots (filled cells), with sole_only consisting of sole crops only, inter_only (sparse) consisting of intercrops only, and sole_inter_50 (sparse) combining both sole crops and intercrops.

**Figure 3 caption:**
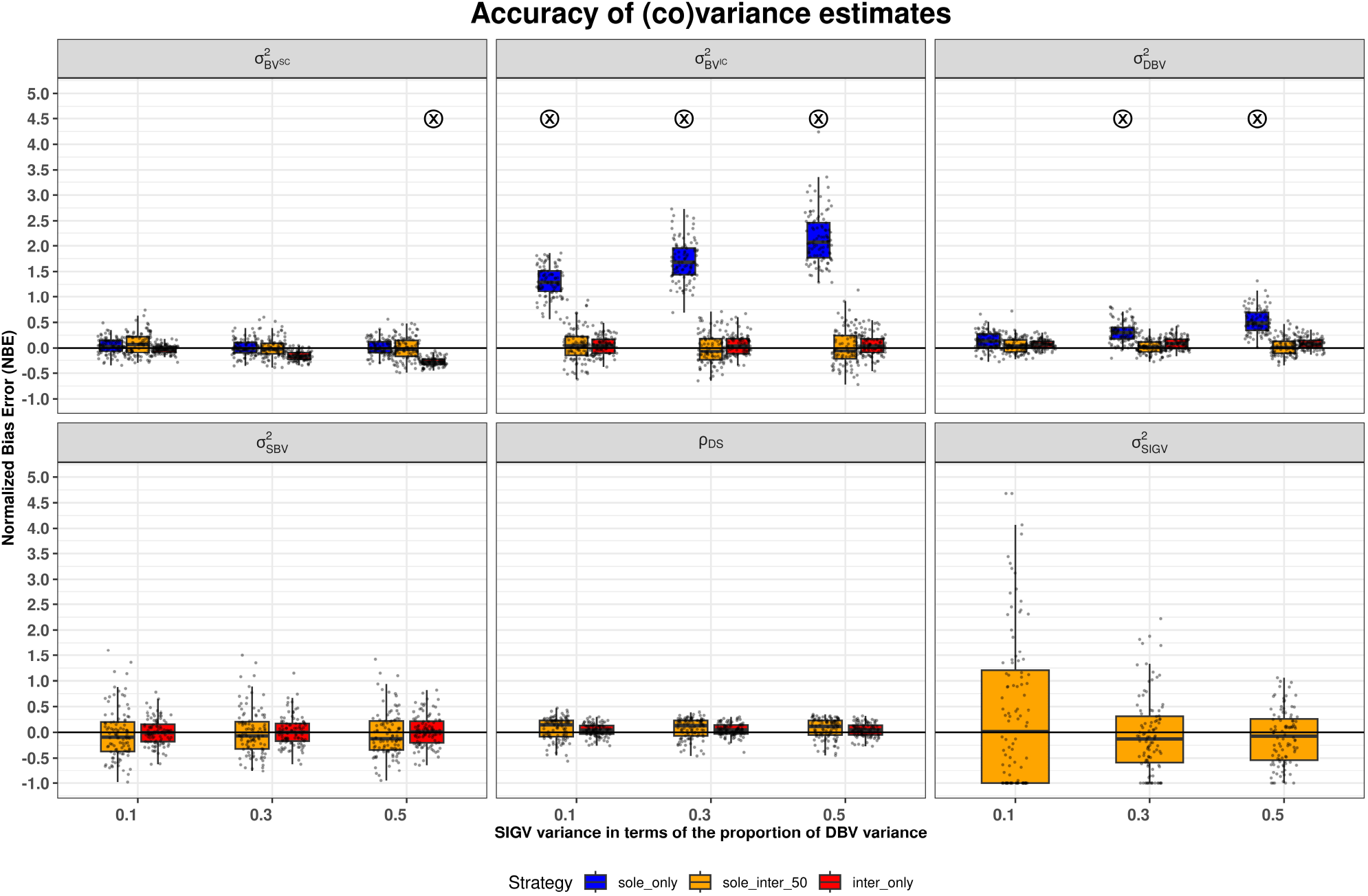
Estimation accuracy (normalized biased error on the y-axis, one dot per simulation) of genetic (co)variances for the focal species, under varying levels of SIGV variance (x-axis) and var(SBV) = 20% var(DBV) . ⓧ marks cases where the bias is significantly greater than 15 % of the mean (|NBE| ≥ 0.15).

**Figure 4 caption:**
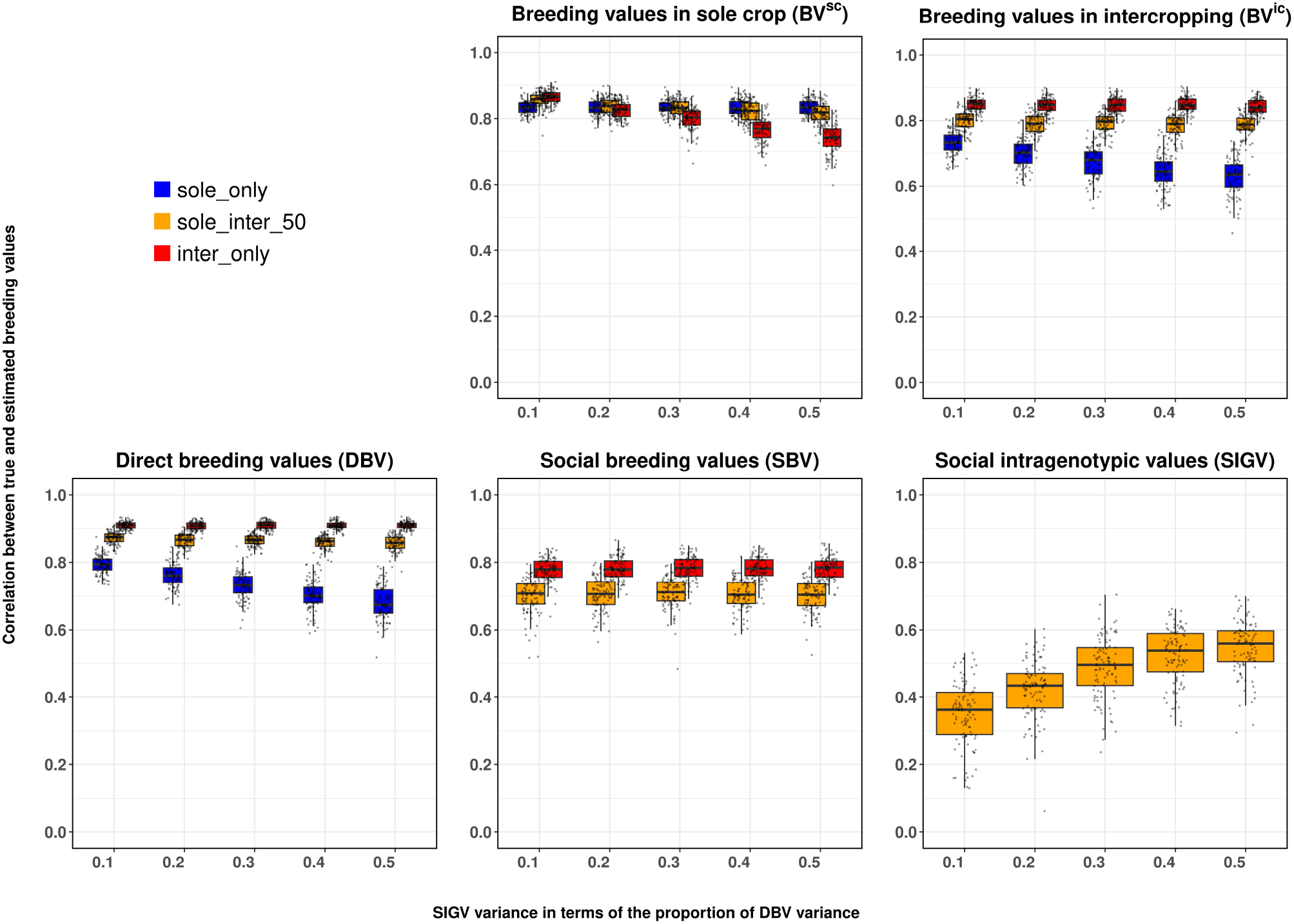
Pearson correlation between true and estimated breeding values for BV^SC^, BV^IC^, DBV, SBV, and SIGV on the y-axis, along scenarios with increasing importance of SIGV, i.e., increasing ratio var(SIGV)/ var(DBV) on the x-axis (var(SBV) = 20% var(DBV), cor(DBV,SBV=-0.8). For the inter_only design, the true classical breeding value was compared with the estimated direct breeding value only, as SIGV was not estimable with this design.

Multiple simulations were replicated for different combinations of parameters (“scenarios”), and estimation accuracy was assessed with the normalized bias error (NBE; the closer to zero, the better). For all genetic variances and correlation of the focal species, over all scenarios, the sole_inter_50 design was the only one without bias larger than 10%, whatever the variance of SIGV (Figure 3). It is also the only one giving access to SIGV variance, although the estimates of this parameter showed the largest variability. Moreover, only the sole_inter_50 and inter_only designs allow estimating the DBV and SBV variances, as well as their covariance. The inter_only design also led to unbiased estimates, except a slight underestimation of the variance of solecrop breeding values, 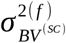 (for SIGV variance >10% of the 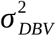), with a bias increasing with SIGV variance. The sole_only design led to large overestimations of 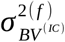, (increasing with SIGV), slight overestimates of the DBV variance, and unbiased estimates of the variance of solecrop breeding values 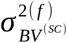. The biases observed are due to the approximations made (eg. for sole_only, 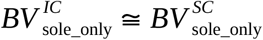 as detailed in the Material and Methods section).

### Accuracy of breeding value estimates

The same simulations were used to assess the accuracy of breeding value estimates (Figure 4). For the sole crop breeding values (BV^SC^), all three designs had a similarly strong accuracy at low SIGV variance. However, with increasing SIGV variance, the sole_only design retained its accuracy, as expected, whereas the sole_inter_50 accuracy was slightly decreasing and inter_only was the most impacted. For the intercrop breeding values (BV^IC^), the inter_only design displayed the highest correlation, stable across all SIGV variances, closely followed by sole_inter_50, also unaffected by the SIGV variance, whereas sole_only was markedly lower and kept getting worse with increasing SIGV variance.

As in the preceding section, these trends were due to the underlying breeding values. The sole_only design showed lower performance, as sole_inter_50 and inter_only exploit the correlation between DBVs and SBVs, resulting in higher accuracy for DBV estimates. The inter_only design produced the most accurate estimates of DBV and SBV, sole_inter_50 being nearly as good, and the only design to provide SIGV estimates.

### Accounting for genetic relationships among individuals

The impact of accounting for genetic relationships (Figure 2 in File S2) on the accuracy of estimated breeding values was assessed in the sole_inter_50 sparse design (Figure 5 and Figure 8 in File S2). Overall, taking genetic relationships into account in the inference increased accuracy of all breeding values, significantly so for the social ones SBVs and SIGVs.

**Figure 5 caption:**
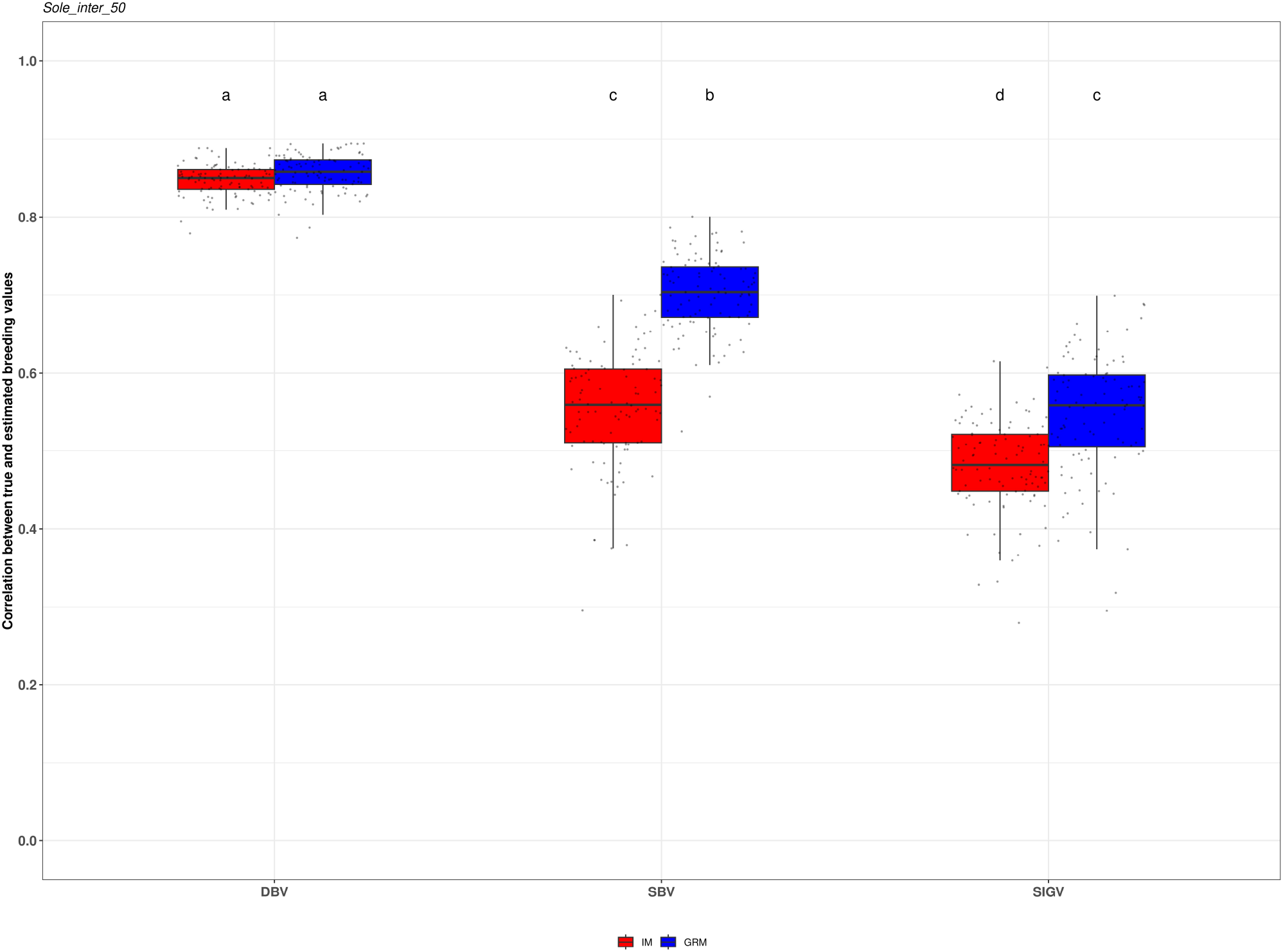
Pearson correlations between true and estimated breeding values (y-axis) for DBV, SBV, and SIGV (x-axis). Parameter relationships are defined as var(SBV)= 20% var(DBV) and var(SIGV) = 50% var(DBV). Estimates obtained with an identity matrix (IM) are shown in red, whereas estimates with a genomic relationship matrix (GRM) are shown in blue. Groups that do not share a common letter differ significantly according to Tukey’s HSD test at the 5% significance level.

### Joint selection for both sole and intercrops

The ability of the three experimental designs to identify the top genotypes for both sole cropping and intercropping was evaluated over a range of SBV and SIGV variances. The true positive rate was defined as the proportion of genotypes selected from estimated breeding values that matched the truly best genotypes (Figure 6 and Figure 9 in File S2).

**Figure 6 caption:**
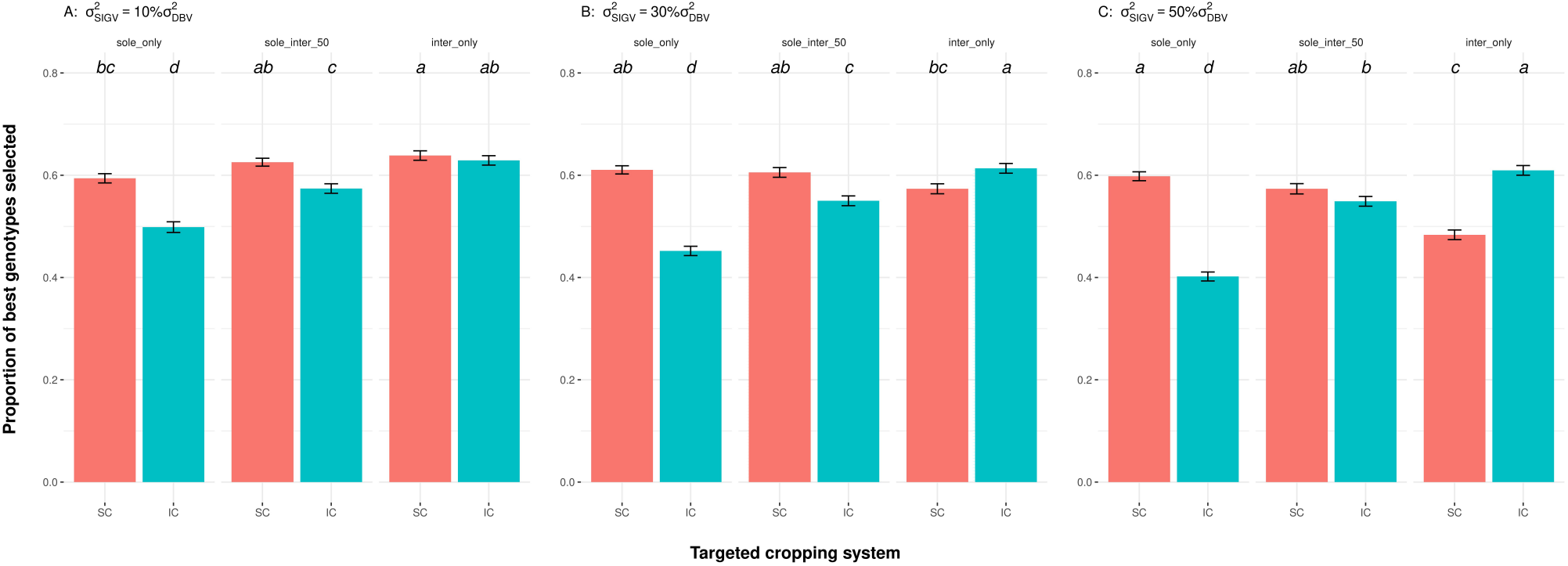
Proportion of top-performing genotypes (y-axis) selected for a targeted cropping system (x-axis; sole cropping, SC, or intercropping, IC), based on their breeding values estimated from three experimental designs (sole_only, sole_inter_50 and inter_only). Results are shown across increasing variance of social intragenotypic values: var(SIGV) = 10% (A), 30% (B), and 50% (C) of var(DBV). Other (co)variance parameters were fixed as follows: var(SBV) = 20% var(DBV), and cor(DBV,SBV) = -0.8. Error bars represent standard errors of the means across replicates. Tukey’s HSD test was performed separately within each level of var(SIGV); groups that do not share a common letter differ significantly (α = 0.05).

The sole_only design showed significantly better performance for breeding for the sole cropping system than for intercropping, with differences becoming increasingly pronounced as SIGV variance increased. The sole_inter_50 design also achieved high accuracy for breeding for sole cropping while maintaining strong performances for intercropping, and was notably less sensitive to increasing SIGV variance. In contrast, the inter_only design was optimal for identifying superior genotypes for intercropping, and also performed well for sole cropping at low SIGV variance (10% of DBV variance). Interestingly, at low SIGV variance, selection for sole crop was significantly better with the inter_only design than with the sole_only design. Indeed, even though the SIGV variance is not captured by inter_only, this was more than compensated by the fact that the DBVs were better estimated with inter_only than sole_only (c.f. Figure 4). At low SIGV variance (10% of DBV variance), neither the sign nor the magnitude of the DBV-SBV correlation change qualitatively the results (Figure 9 in File S2), even though they have a major impact on the total BV^IC^ variance (Figure 10 in File S2).

## Discussion

### A new genetic model of social effects to clarify the correlation between mono- and pluri-specific breeding values

While the question of correlation between solecrop and intercrop genetic values has been highlighted as a critical question (Annicchiarico et al., 2019, Sampoux et al, 2019), the use of both sole crop (SC) and intercrop (IC) performances to increase genetic gain when breeding for intercropping systems has been only recently proposed by Bančič et al., (2021). Indeed, these authors developed genomic predictions by integrating monocrop grain yield as a correlated trait in a multivariate model. However, they did not address how SC and IC breeding values (BV^SC^ and BV^IC^) are interwoven. We hence introduced a unified genetic description of BV^SC^ and BV^IC^ in terms of their underlying genetic determinants: direct and social breeding values (DBV, SBV), and the statistical deviation (DBV×SBV). To account for the difference between BV^SC^ and DBV (that is also part of the BV^IC^), we introduced the social intragenotypic value (SIGV) that cannot be estimated using SC or IC data alone.

The definition of SIGV involves the SBV^SC^ and the (DBV×SBV)^SC^ in the sole crop of the genotype on itself, both terms being indistinguishable in the absence of varietal mixture data. This was motivated by the earlier introduction of intra-genotypic interactions in sole crop when analyzed jointly with varietal mixtures (SMA_ii_) by Forst et al. (2019). Although these authors also proposed to consider such intra-genotypic interactions within varietal mixtures (their model 3), we did not pursue this line here under the assumption that these social intragenotypic effects were likely to be diluted in the intercropping case. However, it remains a perspective to be explored, along with the extension of our framework to analyze monovarietals, cultivar mixtures and species mixtures, jointly.

Additionally, our model decomposes the specific mixing ability in intercrop i-j into two (DBV×SBV)^IC^ terms, i.e., the specific effect of genotype i of species S1 on the yield of genotype j of species S2, and reciprocally. This formulation is made possible by the fact that data are available on both species in intercrops. Compared to the host-pathogen coGWAS case where, typically, only the host disease level is analyzed (M. Wang et al., 2018), our framework could allow the analysis of host disease level and pathogen load jointly. Statistically, we modeled the (DBV×SBV)^IC^ terms as random effects, with one genetic variance per species, a genetic covariance between both species yields, and a genetic similarity matrix between mixtures. We did not explore the influence of these (co)variances in our simulations, notably because further developments are needed to address the genetic similarity between mixtures—a topic we consider beyond the scope of the present paper.

In terms of definition, SBV_.|i_^IC^ is the effect of the ith focal genotype on any tester yield, measured in units of the tester species. Concretely, the SBV_.|i_^IC^ of a wheat genotype i intercropped with pea will be a pea yield, i.e., the specific yield deviation it is responsible for on any pea genotype. In terms of biological interpretation, such SBV^IC^ would relate to biological processes involved in interspecific plant-plant interactions, such as competition, complementarity and facilitation with respect to resources (e.g., Hinsinger et al., 2011) as well as signaling between species (Poveda & Kessler, 2012; N. Wang et al., 2021). Similarly, SIGV relates to intra-genotypic interactions such as density tolerance or architectural traits selected under monoculture conditions, interpreted as cooperative behavior within a population (Denison et al., 2003; Donald, 1968). However, SBV^IC^ and SIGV of the focal genotypes being statistical constructs, their estimated value on any given dataset will be influenced, not only by the specific testers and focal genotypes used in the panel, but also by the relative amount of sole crops vs intercrops in the design. It hence remains an open question to precisely map such genetic values to processes underlying plant-plant interactions, an endeavour out of the scope of the current work.

The main strength of our framework lies in its more detailed decomposition of breeding values BV^SC^ (as DBV+SIGV) and BV^IC^ (as DBV+SBV^IC^) which allows a breeder not only to select for a higher BV^IC^, but also for a balance between both intercropped species. This is particularly important as yield ratios between species can vary strongly according to the outcome of plant-plant interactions between them. With our framework, the breeder hence has the possibility to fine-tune the weights of DBV, SBV^IC^ and SIGV of the focal species in a broader selection index: Idx = w_DBV_DBV + w_SBV_SBV^IC^ + w_SIGV_SIGV. Moreover, in an ideotyping approach, our framework also allows a breeder to identify traits more specifically related to SBV^IC^ than DBV or SIGV for instance, hence providing a better understanding of the genetics of intra- and inter-specific interactions. Furthermore, beyond breeding applications, our decompositions provide access to genetic values such as SBV^IC^ and SIGV that are worth studying for themselves, e.g., to investigate their genetic architecture, their dynamics across generations (notably the covariance between DBV and SBV), whether through simulations or experimental studies.

### An efficient design with both sole and intercrops

Breeding for mixtures suffer from the combinatorial explosion of the number of possible assemblages. We hence developed a strategy based on the joint analysis of monovarietals and mixtures to allow breeders to gradually incorporate mixing ability in their breeding target. Our strategy is based on incomplete designs and our results here are consistent with the findings of Haug et al. (2021), who demonstrated the efficiency of incomplete designs for estimating genetic (co)variance parameters in the context of intercropping breeding.

Among the three evaluated designs, the joint analysis (sole_inter_50) was the only one that enabled estimation of all genetic (co)variances, and it showed strong performance compared to the sole_only design. As expected, each design showed advantages for specific breeding values: sole_only provided the most accurate estimates of BV^SC^, while inter_only yielded the best estimates of BV^IC^. The large bias from sole_only design when approximating BV^IC^ arises from the fact that: i) the sole_only design cannot estimate social breeding values (SBV) due to the absence of companion species; ii) it cannot exploit the DBV-SBV correlation, that contributes, when present, to higher accuracy; iii) the presence of SIGV in the sole_only design——further inflates the bias in BV^IC^ approximation.

Interestingly, at low SIGV, the sole_inter_50 and inter_only designs were better at identifying the best genotypes than the sole_only design. This improved performance is likely due to the greater amount of data available in the designs with intercrops. However, this additional phenotyping—effectively corresponding to an extra trait—comes at a cost (e.g., due to the need to sort grains per crop after harvest) that requires careful consideration when planning the breeding design, even when the total number of plots remains unchanged. Indeed, grain sorting requires a specific machinery (Fouillet et al., 2025) and hence has a cost for both farmers and breeders, but no specific study addressed the issue for the latter. Previous work that compared alternative breeding strategies, such as phenotypic vs genomic prediction (e.g., Marulanda et al., 2016), considered a unique cost for a field plot. Therefore such approaches will have to be extended to the intercropping case in this regard.

Even though we only investigated tester-based designs here, motivated by the results of field trials (Moutier et al., 2022), the question remains to compare them with incomplete factorials. Genomic prediction is likely to be a key ingredient in such comparisons, as was done for hybrid breeding (Seye et al., 2020). Our model will also be useful in such cases as it allows to leverage a genomic relationship matrix, and could be easily extended to accommodate any kernel-based matrix as is done in classical ridge regression (Endelman, 2011). Moreover, future extensions incorporating kernel-based, crop model-based and/or deep learning approaches could also be leveraged in order to capture the non-linear processes underlying plant development and plant-plant interactions. One possibility in this direction would be to combine a crop model able to simulate both sole crops and intercrops (e.g., Vezy et al., 2023) with a deep learning approach, as was done by Han et al., (2025) for monovarietal conditions.

### Relative weighting of sole-crop and/or intercrop breeding targets

The relevance of sole-crop information for intercrop breeding depends on the correlation between BV^IC^ and BV^SC^, as well as on the heritabilities h^2^_SC_ and h^2^_IC_ . In our simulation scenarios, the heritabilities were assumed to be equal in both sole crops and intercrops. Therefore, the relative efficiency of indirect selection in sole crops versus direct selection in intercrops depended solely on the correlation, which ranged from 0.92 to 0.58 (Figure 11 in File S2). These results indicate, first, that indirect selection in sole crops was always less efficient than direct selection for intercrops; and second, that as social interactions intensify (i.e., with increasing SIGV and SBV), this inefficiency increases.

The range of correlation considered here included the values observed on barley-pea mixtures in four environments (Haug et al., 2023) and on wheat–pea mixtures in three environments (Salomon et al., in prep.). Other authors explored even lower correlations, e.g., as low as 0.4 in Bančič et al., (2021). Reaching such values would require even larger SIGV and SBV variances, as well as including non-zero DBV×SBV variances. In our simulations, interspecific interactions captured by the statistical deviation (DBV× SBV)^IC^, were not considered as, in published datasets of barley-pea mixtures, these interactions were negligible (Haug et al., 2023). This may be explained by a small phenotypic contrast between testers as well as their limited number, for practical and financial reasons. However, such interactions may be important in other, specific biological contexts and deserve further investigation.

In the current context, where sole-crop systems dominate market share, it appears more pragmatic to gradually integrate intercrop evaluation into solecrop breeding programs rather than to opt for a drastic shift toward dedicated intercrop breeding programs. In this regard, the sole_inter_50 design, which combines sparse and joint evaluation under both solecrop and intercrop conditions, represents an effective strategy for breeding programs seeking to allocate part of their resources to intercrop evaluation. The specific value of 50% of mixtures used in this study can then be adjusted depending on the breeding goals and on the market. The weight *w*_*sole*_ in the new selection index for this sole_inter design can also be adjusted according to these aspects. Furthermore, accounting for the specific economic values is critical for breeders, as intercropping can involve species with contrasted values (Vandermeer, 1989). Our framework straightforwardly allows this when applied directly on the specific economic value rather than the yields. Another approach would be to fine-tune the weights of the broader selection index mentioned above (w_DBV_, w_SBV_ and w_SIGV_) with respect to the specific economic values.

The joint strategy proposed in this study (in terms of genetic and statistical modeling as well as experimental design) could also be effective for multi-generation selection in both species and cropping systems, as they would allow a large number of genotypes to be tested. When combined with genomic selection (Danguy des Déserts et al., 2023; He et al., 2016), these strategies could help achieve high genetic gain at lower cost compared with complete full factorial designs. Further simulation studies using true parameters should be conducted to identify at which stage(s) of breeding schemes these approaches are more efficient. To achieve a complete view, the impact of early selection among contrasted ideotypes, using trait combinations correlating with social breeding values should also be considered in parallel to a more quantitative genetics approach as studied here.

## Conclusion

Our results highlight the efficiency of a new genetic and statistical framework to jointly evaluate mono- and pluri-specific performances in incomplete designs. This approach supports breeding strategies where breeders want to gradually integrate pluri-specific trials in their mono-specific platforms. We showed that the simultaneous improvement of breeding values in mono- and pluri-specifics crops is reachable, mono-specific trials providing information on the direct breeding values also present in pluri-specifics and, reciprocally, pluri-specifics providing valuable insights into the mono-specific performance. The efficiency and necessity of developing specific breeding design for pluri-specifics is strongly relying on the importance of the variance on social breeding values either in mono- or pluri-specifics, which in turns depends on the studied species, and their management (such as sowing density and fertilization for crops). Therefore, translating these theoretical findings into practical breeding recommendations requires experiments to assess these variance components and accordingly optimize breeding schemes integrating, the extra cost of evaluating pluri-specific trials. Beyond multi-specific groups in crops, our framework paves the way to the genetic study of such groups in animals (Goodale et al, 2020) and trees (Tang et al, 2025) up to whole ecosystems (Whitman et al, 2006).

## Data availability statement

All the code and data to reproduce the results is available in the Research Data Gouv Repository, at https://doi.org/10.57745/YU9T9X. We also provided a R package, **plantmix** (https://cran.r-project.org/package=plantmix), which implements the methods developed in this study and offers additional functionalities.

## Acknowledgments

The authors want to thank the members of the MoBiDiv project for fruitful feedback, notably J. David and G. Montazeaud.

## Author contributions

All authors established the genetic model, statistical framework and simulation plan. JS implemented them in R and TMB, supervised by TF. All authors analyzed the results and reviewed the final manuscript.

## Supplementary information

The supplementary material for this article can be found online inin the Research Data Gouv Repository, at https://doi.org/10.57745/YU9T9X.

