## Supplemental Material 1 for "Joint modeling of social genetic effects in mono- and pluri-specific groups: case study in intercrops"

**Supplementary File 1**

### 1 Simulations of field trials

#### 1.1 Default parameter values

- Mean yield of wheat genotypes (in q/ha): $\mu^{(f)} = 65$ in sole crops, and reduced by half in intercrops ($65 \times 0.5$)

- Mean yield of pea genotypes (in qt/ha): $\mu^{(t)} = 30$

- Wheat DBV variance: $\sigma_D^{2(f)} = (CV_g^{(f)} \times \mu^{(f)})^2$ where $CV_g^{(f)} \in \{8\%, 8.5\%\}$

- Prop. of wheat $SBV^{(f,IC)}$ variance: $\sigma_S^{2(f,IC)} = 0.2\,\sigma_D^{2(f)}$

- Prop. of wheat SIGV variance: $\sigma_{SIGV}^{2(f)} = 0.5\,\sigma_D^{2(f)}$

- Correlation between wheat DBV and $SBV^{(f,IC)}$: $\rho_{DS}^{(f)} = -0.8$
  - Covariance: $\sigma_{DS}^{(f)} = \rho_{DS}^{(f)} \;/\; (\sigma_D^{(f)} \times \sigma_S^{(f)})$

- Variance of $(DBV \times SBV)^{IC}$: $\sigma_{D\times S}^{2(f)}(\text{resp.}, \sigma_{D\times S}^{2(t)}) = 0$

- Variance of weat $BV^{SC}$: $\sigma_{BV^{SC}}^{2(f)} = \sigma_D^{2(f)} + \sigma_{SIGV}^{2(f)}$

- Variance of wheat $BV^{IC}$: $\sigma_{BV^{IC}}^{2(f)} = \sigma_D^{2(f)} + 2\,\sigma_{DS}^{(f)} + \sigma_S^{2(f,IC)}$

- Sole error variance for focal: $\sigma_{e,sc}^{2(f)} = \frac{1-h_{sc}^{2(f)}}{h_{sc}^{2(f)}} \times (\sigma_{BV^{SC}}^{2(f)} \times r)$ where $h_{sc}^{2(f)} = 0.7$, and $r$, the number of replicates (2)

- Intercrop error variance was approxiamte for both, tester and focal by: $\sigma_{e,ic}^2 = \frac{1-h_{ic}^2}{h_{ic}^2} \times (\sigma_{BV^{IC}}^2 \times (t \times r))$ where $h_{ic}^2 = 0.7$, $t$ the numer of effective tester, and $r$ the number of replicates

- Correlation between mixed focal and tester errors: $\rho_{e,ic}^{(f,t)} = -0.2$
