## Supplemental Material 2 for "Joint modeling of social genetic effects in mono- and pluri-specific groups: case study in intercrops"

### Supplementary File 2

**Supplementary Figure 1:** Linkage disequilibrium between SNPs in the simulated data set

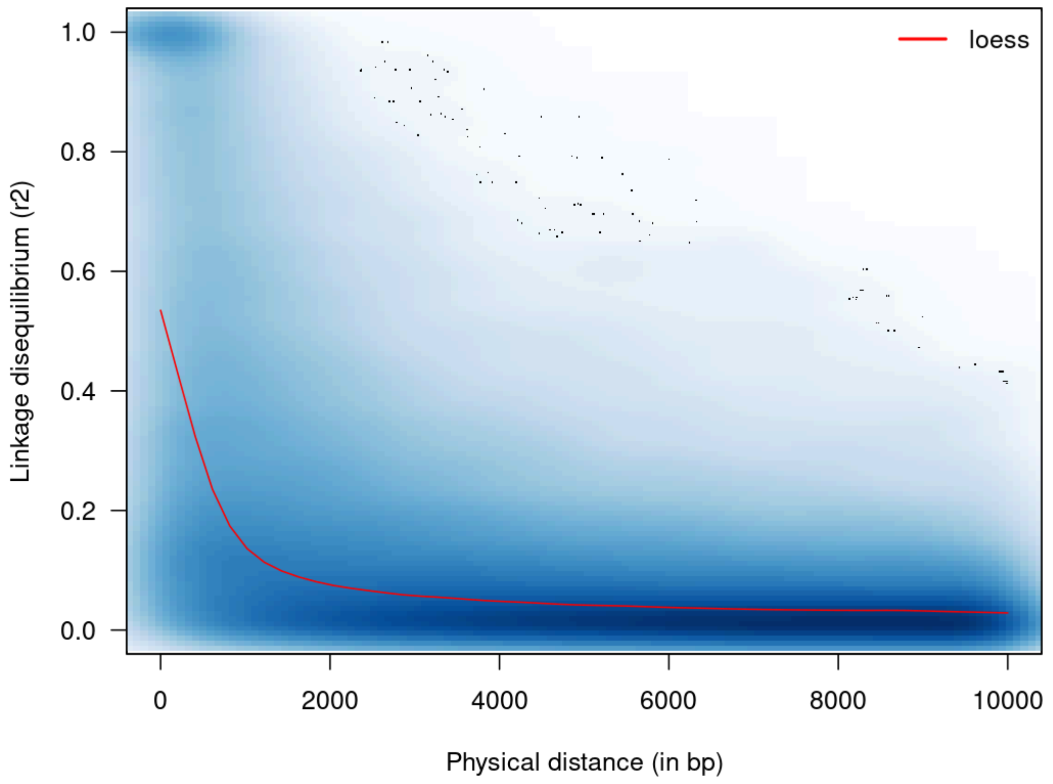

**Supplementary Figure 2:** Genomic relationship matrix in the simulated data set

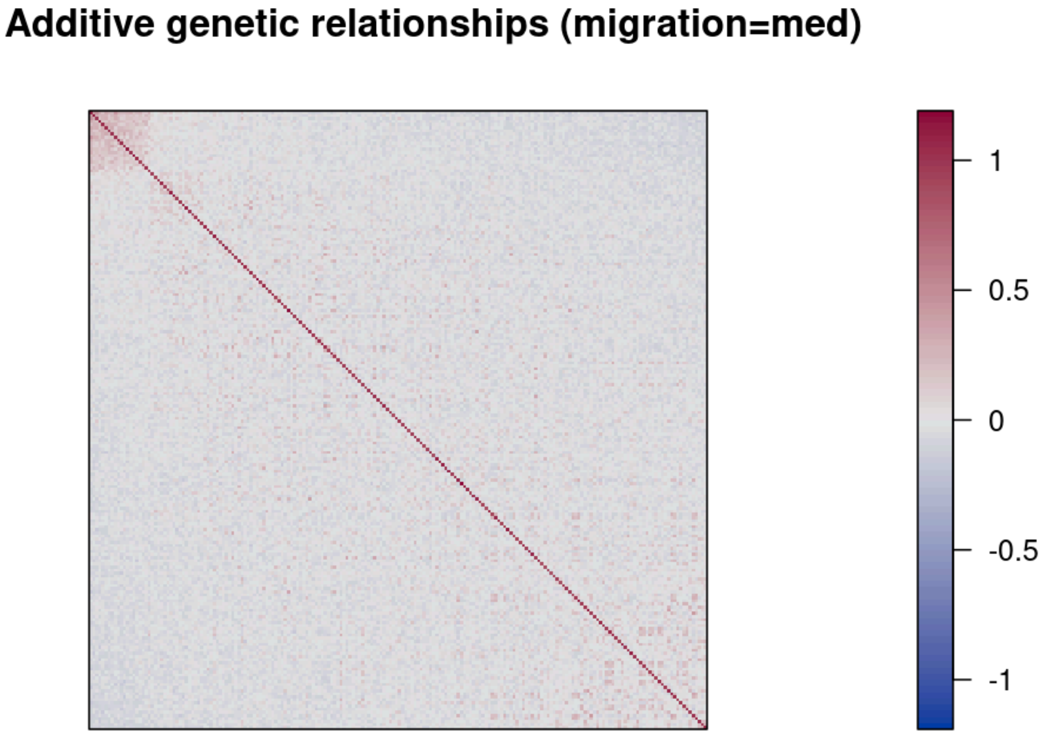

**Supplementary Figure 3:** Genetic structure among genotypes in the simulated data set

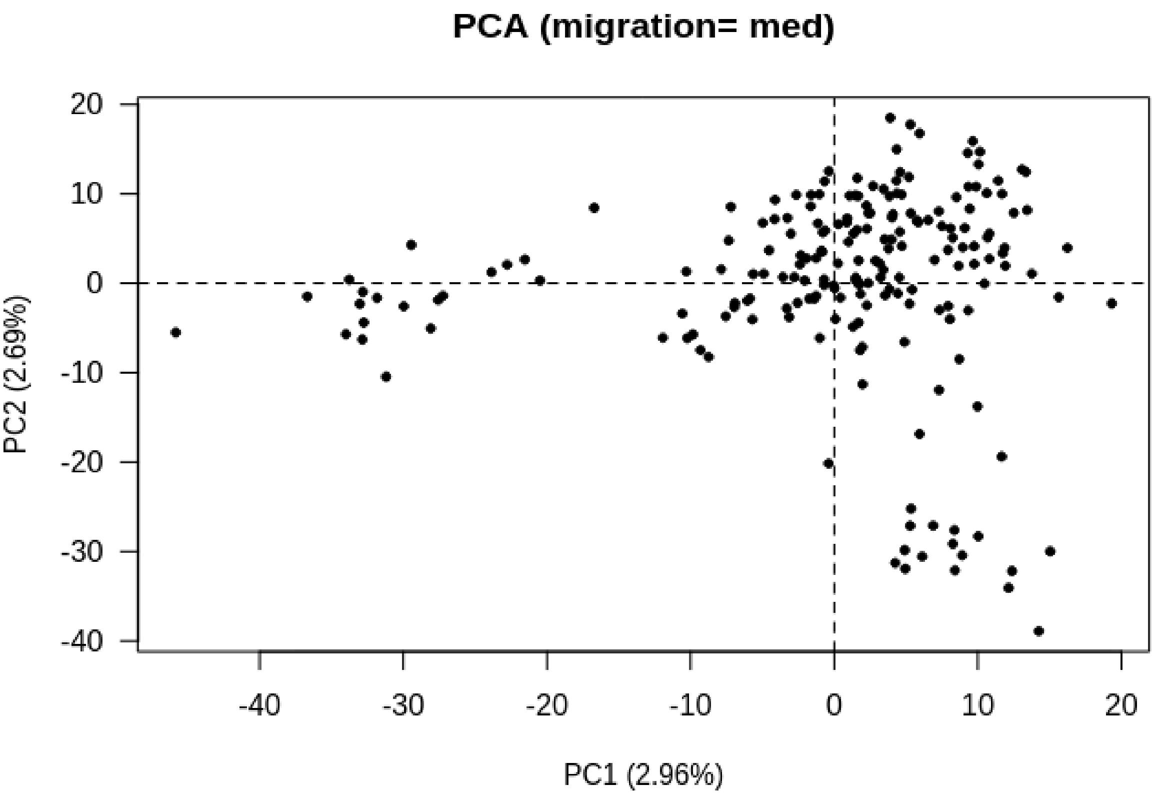

**Supplementary Table 1:**Computational efficiency of ASReml and TMB

| **Model** | **Designs** | **TMB**<br><br>**(in seconds)** | **ASReml-R**<br><br>**(in seconds)** |
|---|---|---|---|
| *Univariate* | `sole_only` | *Auto. diff.: 37.00*<br><br>*Nll[1] minimisation (R): 4.30*<br><br>*Total: 41.30* | *Total: 0.10* |
| *Multivariate* | `inter_only` | *Auto. diff. : 30.38*<br><br>*Nll[1] minimisation (R): 27.15*<br><br>*Total: 57.53* | *Total: 0.27* |
| | `sole_inter_50` | *Auto. diff.: 16.05*<br><br>*Nll[1] minimisation (R): 166.01*<br><br>*Total: 172.06* | *NA* |

[1]: negative log-likelihood

**Supplementary Table 1:** The computational efficiencies of ASReml-R and TMB were assessed for each design. The values in the table correspond to averages over 100 replicates, simulated with var(SBV) = 20% var(DBV), var(SIGV) = 10% var(DBV) and cor(DBV, SBV) = -0.8.

**Supplementary Figure 4:** Estimation accuracy of ASReml and TMB

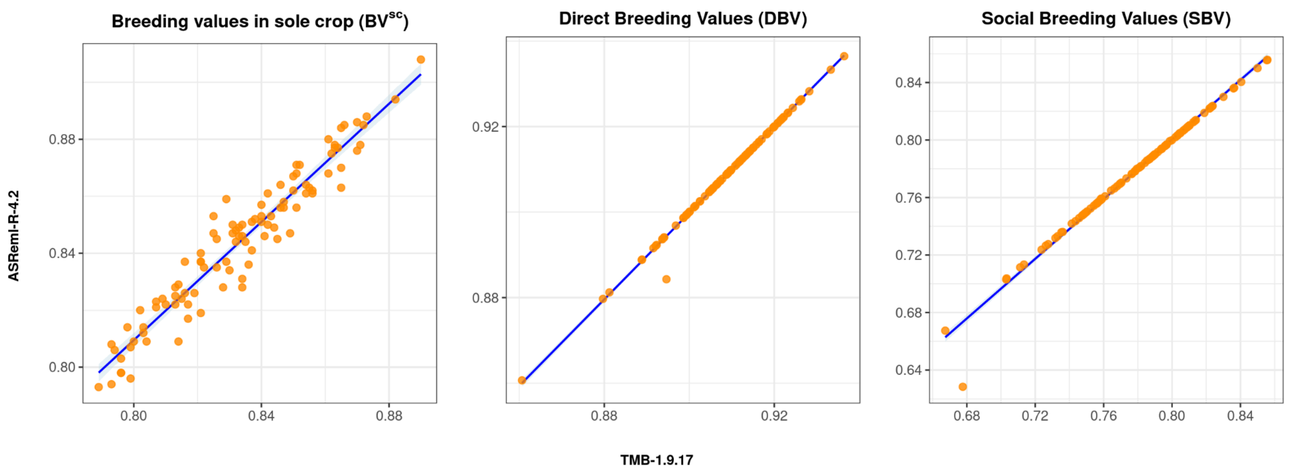


**Supplementary Figure 4:** Comparison of estimated breeding values obtained using ASReml-R 4.2 and TMB 1.9.17. The figure presents breeding values in solecropping ($BV^{SC}$) estimated from the sole_only design, the direct breeding values (DBV) and social breeding values (SBV) from the inter_only design. Each panel displays a scatter plot comparing estimates from both methods across replicates, with individual points representing data distributions.

TMB and ASReml-R exhibited an almost perfect correlation for the estimation of direct breeding values (DBV: Radj2 = 0.992, RSE=0.001) and social breeding values (SBV: Radj2 = 0.984, RSE=0.005), but a slightly less perfect correlation for the breeding values in solecrop ($BV^{SC}$: Radj2 = 0.924, RSE=0.007).

**Supplementary Figure 5:** Selection of genotypes in a bivariate distribution

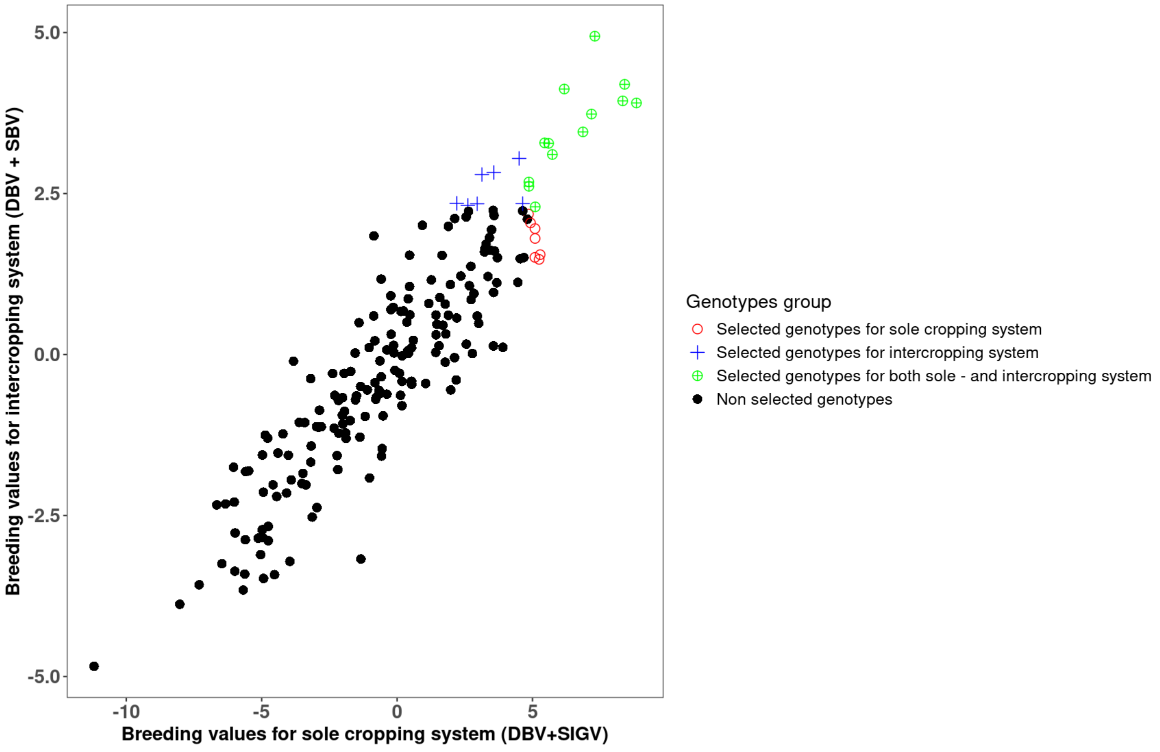


**Supplementary Figure 5:** Selection of genotypes in a bivariate distribution on one simulated dataset with the selection index for the sole cropping system ($BV^{SC}$=DBV + SIGV) on the x-axis and the selection index for the intercropping system ($BV^{IC}$ = DBV + SBV) on the y-axis. Simulation parameters are defined as follow: sole_inter_50; var(SBV) = 20%var(DBV); var(SIGV) = 50%var(DBV), cor(DBV, SBV)= -0.8. A selection proportion of 10% was first applied independently for the sole cropping system and then for the intercropping system. Genotypes selected for both systems correspond to the common genotypes between them.

**Supplementary Figure 6:** Accuracy of error (co)variance estimates

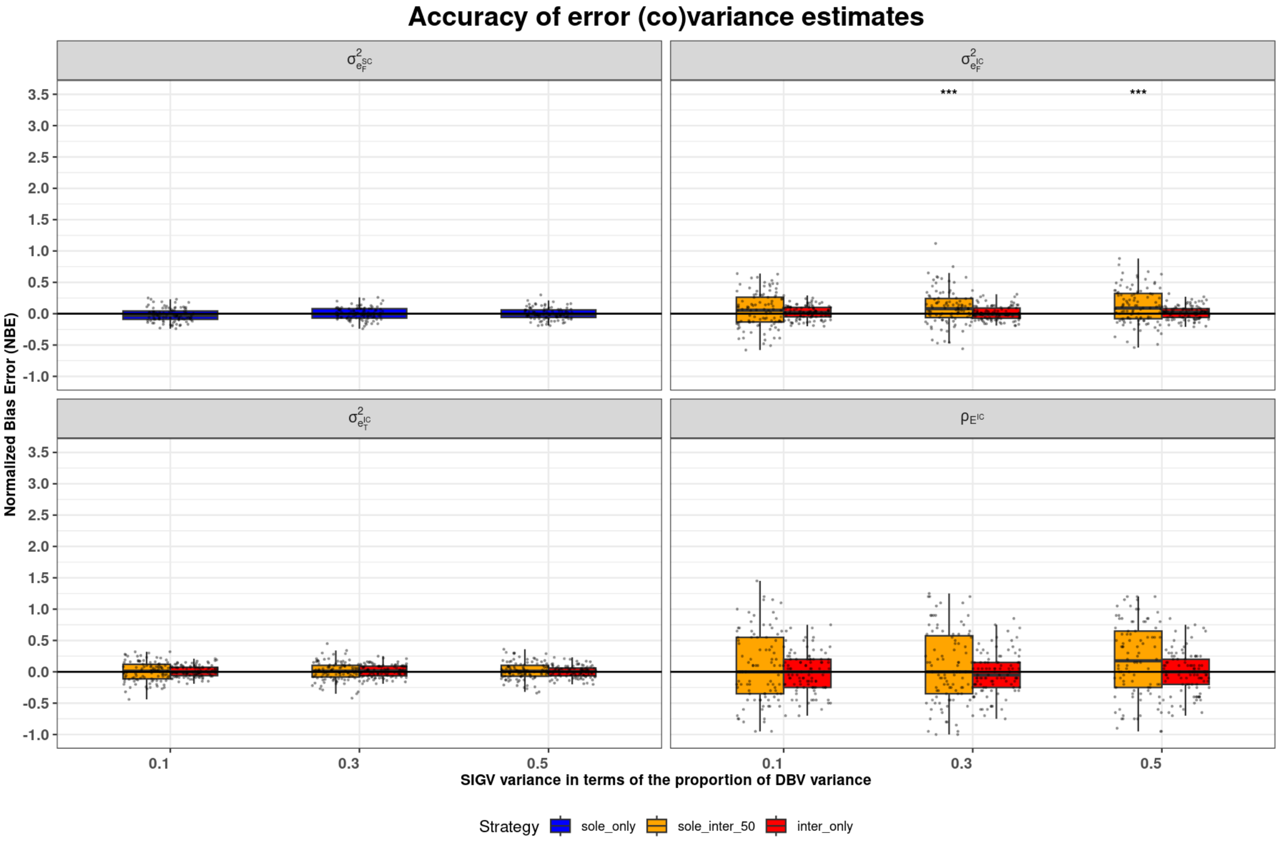


**Supplementary Figure 6:** Estimation accuracy (normalized biased error on the y-axis, one dot per simulation) of error (co)variances(F= focal species; T= tester species), under varying levels of SIGV variance (x-axis)(var(SBV) = 20%var(DBV)). "*" above each box indicate whether the mean normalized biased error is significantly different from zero based on a one-sample t-test: * $p < 0.05$, ** $p < 0.01$, *** $p < 0.001$. Non-significant results are left unlabeled.

**Supplementary Figure 7**: Influence of the social intragenotypic variance on genotypes selection

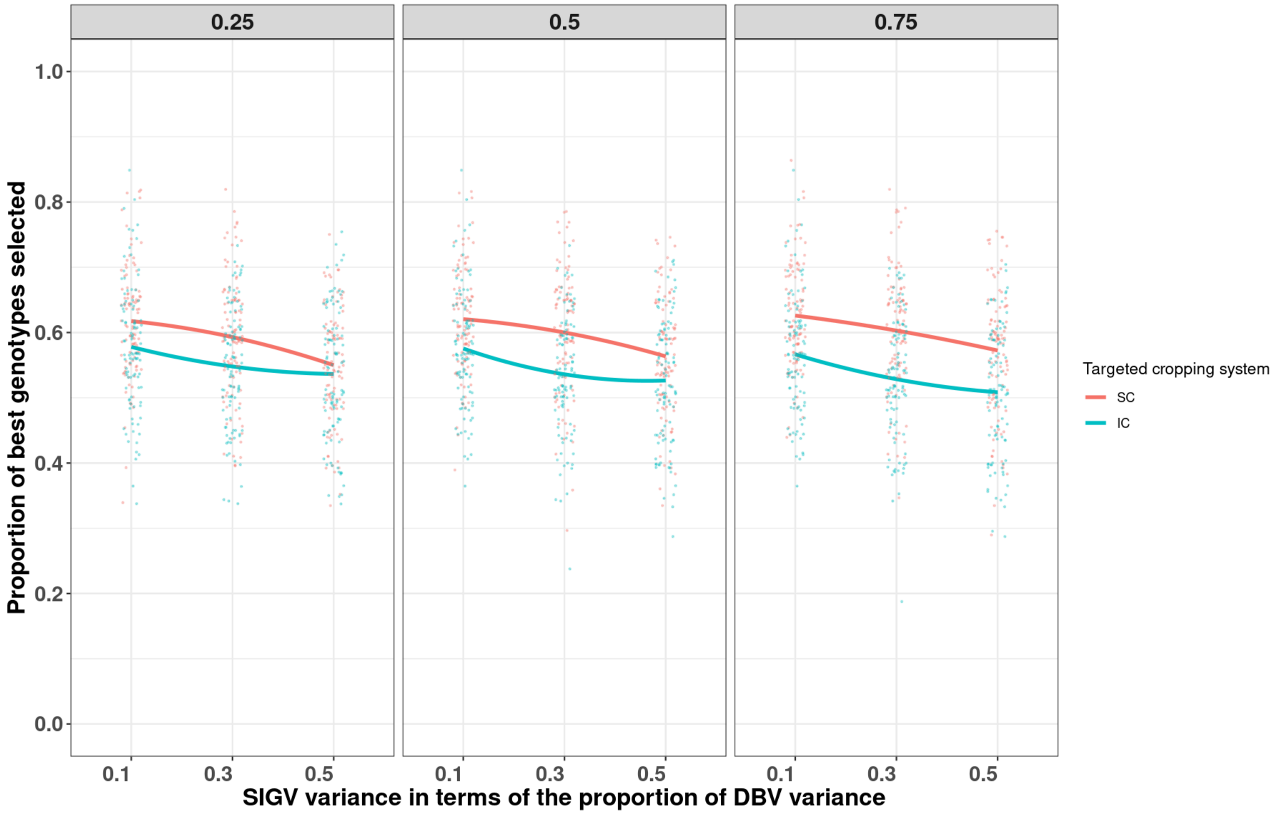


**Supplementary Figure 7:** Proportion of top-performing genotypes selected for sole cropping and intercropping across varying levels of social intragenotypic variance (x-axis), the data being simulated with var(SBV) = 20% var(DBV). Results are shown for $w_{sole}$ = 0.25, 0.50, and 0.75.

**Supplementary Figure 8:** Accuracy with or without genetic relationships over a range of SIGV variance

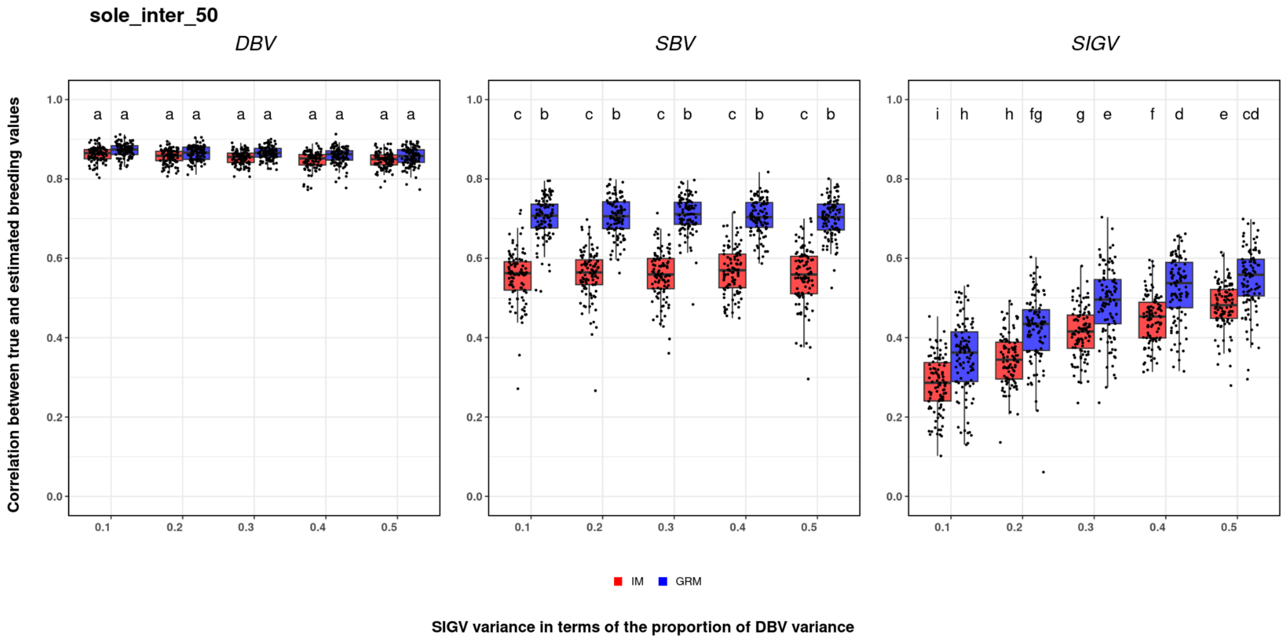


**Supplementary Figure 8**: Pearson correlations between true and estimated breeding values (y-axis) across varying levels of SIGV variance (var(SBV) = 20% var(DBV)). Estimates obtained with an identity matrix (IM) are shown in red, whereas estimates with a genomic relationship matrix (GRM) are shown in blue. Groups that do not share a common letter differ significantly according to Tukey's HSD test at the 5% significance level.

**Supplementary Figure 9**: Influence of the correlation between DBV and SBV on genotype selection

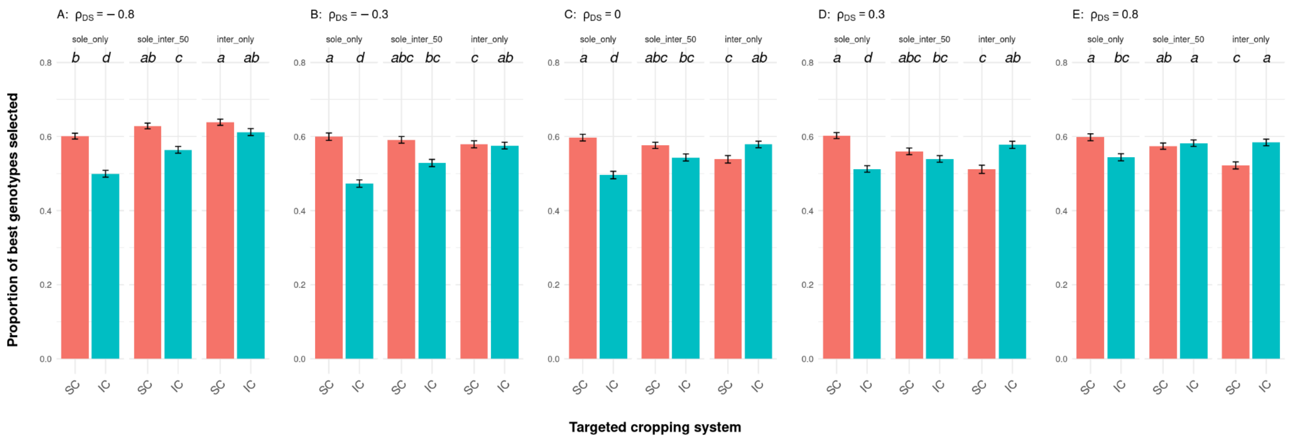


**Supplementary Figure 9**: Proportion of top-performing genotypes (y-axis) selected for a targeted cropping system (x-axis; sole cropping, SC, or intercropping, IC), based on their breeding values estimated from three experimental designs (sole_only, sole_inter_50 and inter_only). Results are shown across varying signs and magnitudes of cor(DBV, SBV): -0.8 (A), -0.3 (B), 0 (C), 0.3 (D) and 0.8 (E). Other (co)variance parameters were fixed as in figure 6: var(SBV) = 20% var(DBV) and var(SIGV) = 10% var(DBV). Error bars represent standard errors of the mean across replicates. Tukey's HSD test was performed separately within each level of cor(DBV, SBV); groups that do not share a common letter differ significantly ($\alpha = 0.05$).

**Supplementary Figure 10:** Influence of the correlation between DBV and SBV on total genetic variance

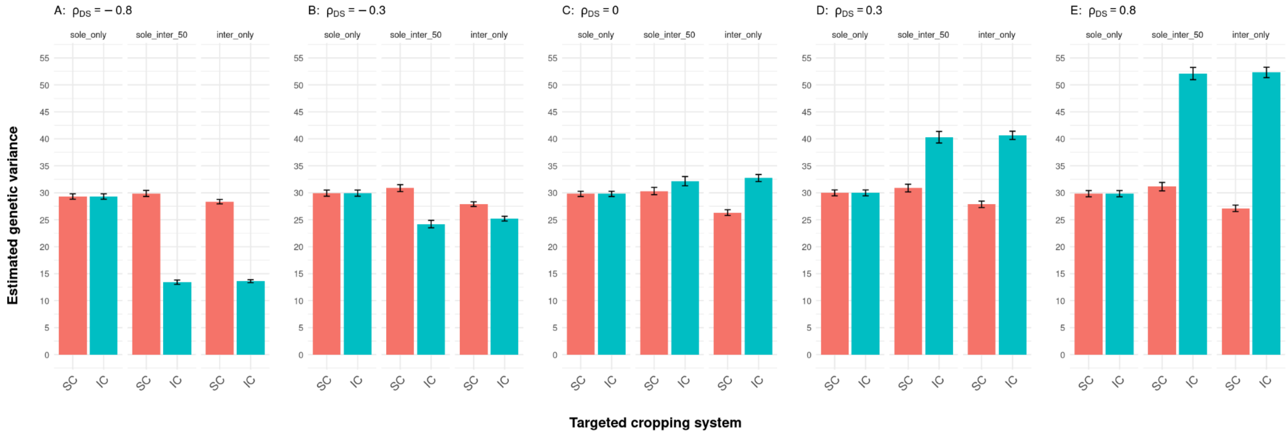


**Supplementary Figure 10**: Estimated genetic variance (y-axis) per targeted cropping system (x-axis; sole cropping, SC, or intercropping, IC), across the three experimental designs (sole_only, sole_inter_50 and inter_only). Results are shown across varying signs and magnitudes of cor(DBV, SBV): -0.8 (A), -0.3 (B), 0 (C), 0.3 (D) and 0.8 (E). Other (co)variance parameters were fixed as in figure 6: var(SBV) = 20% var(DBV) and var(SIGV) = 10% var(DBV). Error bars represent standard errors of the mean across replicates.

**Supplementary Figure 11:** Influence of the social genetic variance on genotypes selection

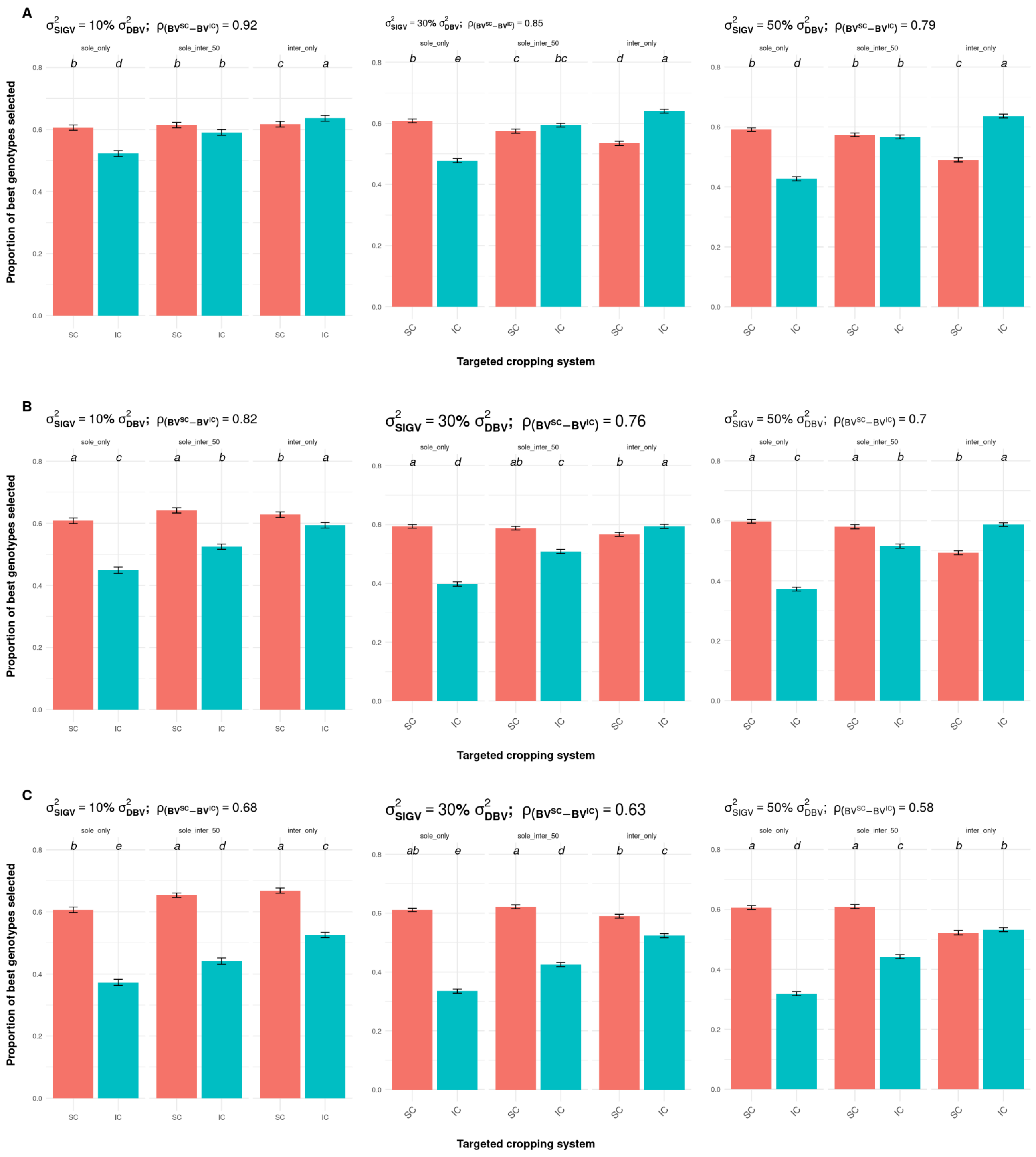


**Supplementary Figure 11**: Proportion of top-performing genotypes selected for sole cropping and intercropping systems across varying levels of social intragenotypic value variance and $BV^{SC}$-$BV^{IC}$ correlation. Results are shown for different levels of SBV variance expressed as a proportion of DBV variance (**A** = 10%; **B** = 30%; **C** = 50%), cor(DBV, SBV) being fixed at -0.8. Error bars represent standard errors across replicates. Tukey's HSD test was performed separately within each SIGV variance proportion; groups that do not share a common letter differ significantly (α = 0.05). The x-axis indicates the cropping system under which selection was applied, where "sole" refers to selection for a sole cropping system and "mix" to selection for an intercropping system. Within each $\sigma^2_{SIGV}$ scenario, three selection strategies—sole_only, inter_only, and sole_inter_50—are compared.

**Supplementary Figure 12:** Accuracy of breeding values with $(DBV\times SBV)^{IC}$

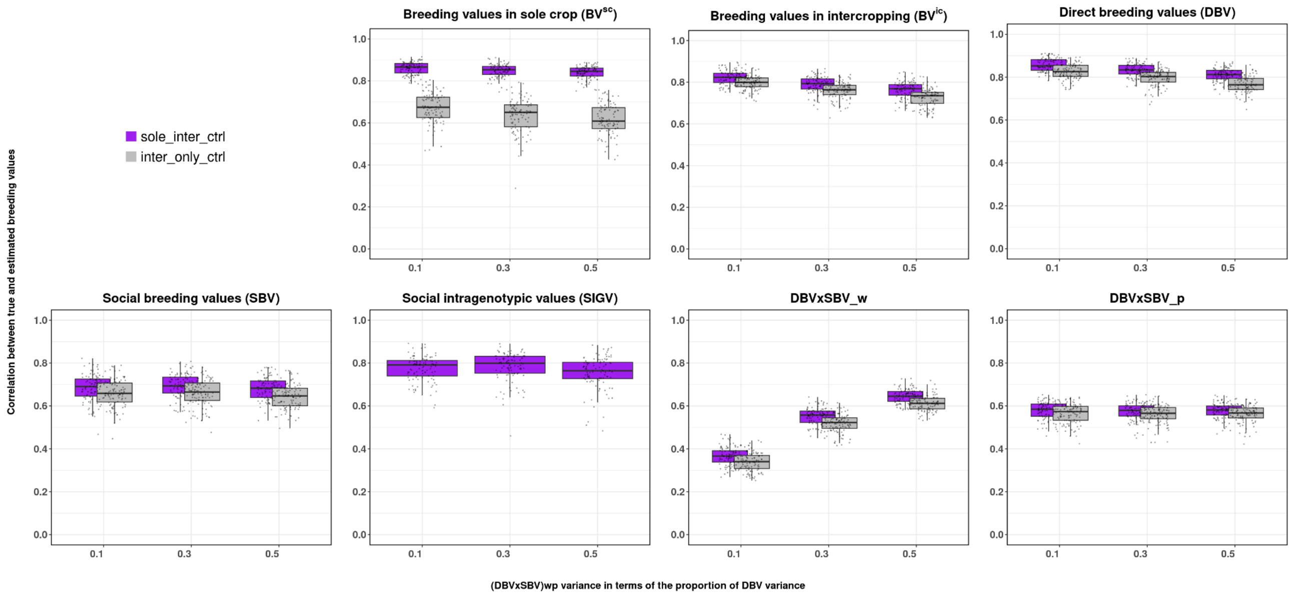


**Supplementary Figure 12**: Pearson correlations between true and estimated breeding values for $BV^{SC}$, $BV^{IC}$, DBV, SBV, SIGV, interaction effects (DBV×SBV_w, which is the $(DBV\times SBV)^{(f|t)}$ ) on focal (wheat) performance and interaction effects (DBV×SBV_p, which is the $(DBV\times SBV)^{(t|f)}$) on tester (pea) performance plotted on the y-axis. Simulation scenarios with increasing importance of the DBV×SBV_w are on the x-axis. DBV×SBV_w is expressed as the ratio var(DBV×SBV_w)/var(DBV). Simulation parameters were var(SBV) = 20%var(DBV), var(SIGV) = 50%var(DBV), cor(DBV, SBV) = −0.8, cor(DBV×SBV_w, DBV×SBV_ p) = 0, and var(DBV×SBV_ p) = 30%var(DBV_ p). For the inter_only design, the true breeding value on solecrop ($BV^{SC}$ ) was compared only with the estimated direct breeding value because SIGV was not estimable under this design. The genetic similarity between mixtures is assumed to be identity $K^{(fxt,\ IC)}$ = Id.
